# Host genetics and diet jointly shape the microbiome of *Drosophila melanogaster* but do not predict lifespan or age-related traits

**DOI:** 10.1101/2025.10.21.683603

**Authors:** Nikolaj Klausholt Bak, Stine Karstenskov Østergaard, Patrick Skov Schacksen, Jeppe Lund Nielsen, Palle Duun Rohde, Torsten Nygaard Kristensen

## Abstract

The microbiome is a key determinant of organismal health, yet inter-individual variability and heterogeneous responses to environmental conditions complicates the understanding of its effects on hosts. Here, we present a comprehensive analysis using the *Drosophila* Genetic Reference Panel (DGRP) to investigate how the interplay between host genetic variation and diet influences microbiome composition, and to assess whether microbiome features in young flies can be used to predict lifespan and age-related traits. Our findings show that adult flies reared on a nutritionally rich control diet exhibited higher microbial richness but lower evenness compared to those on a nutritionally poor restricted diet. Principal component analysis (PCA) highlighted substantial diversity among lines reared on the same diet, and this variation was heritable supported by high heritability estimates for all measured α-diversity metrics, including Unique OTU counts, Shannon and Simpson indices, as well as the relative abundances of genera and species with relative abundances exceeding 1%. These results underscore the critical roles of both environmental factors and genetic variation in shaping microbiome composition under different dietary conditions. Moreover, we identified widespread genotype-by-diet interactions, suggesting that the genetic regulation of the microbiome is highly complex. Finally, we found that the microbiome features of young flies including diversity indices, taxonomic abundances, or ordination scores cannot predict age-associated phenotypes (lifespan, locomotor activity, dry weight, and heat knockdown time). Our findings offer valuable insights into the genetic architecture that governs microbiome composition, dietary responses, and aging in *Drosophila melanogaster*.

**Author Summary:** The community of microbes living on or inside animals - the microbiome - plays an important role in health and aging, but its composition varies greatly between individuals and across environments, making it challenging to fully understand its importance. In this study, we examined how host genetics and diet shape the microbiome of the fruit fly, *Drosophila melanogaster*, using a large panel of genetically diverse lines. We found that the microbiome of flies on a restricted nutritionally poor diet were relatively evenly distributed, while the microbiome of flies on a nutritionally rich diet had a higher species diversity. Many features of the microbiome were strongly influenced by the flies’ genes, and these genetic effects often depended on diet. Finally, we tested whether microbiome features obtained from young flies could predict age-related traits such as lifespan and activity at similar aged flies or later in life, but found no evidence for such links. These results highlight how genetic variation and diet interact to shape the microbiome and its role in aging.

## Introduction

Among the many factors shaping the microbiome, diet is one of the most influential. Both short- and long-term dietary patterns can significantly alter microbial community composition and function [1,2]. In *Drosophila melanogaster*, diet-microbiome interactions are critical, especially under conditions of nutrient scarcity. Commensal bacteria such as *Acetobacter* and *Lactobacillus* enhance larval growth by improving nutrient absorption, stimulating digestive enzyme activity, and synthesizing essential nutrients [3,4]. These effects are particularly pronounced under malnourished conditions, where microbes can serve as a direct protein source, extending lifespan and supporting development [5]. Notably, the impact of microbes on host physiology is highly context-dependent: while beneficial under nutrient-poor conditions, the same microbes may have neutral or even deleterious effects in nutrient-rich environments, depending on their metabolic activity and interactions with the host immune system [5].

The microbiome also plays a significant role in shaping host lifespan and aging. Age-associated changes in microbiome composition and abundance have been consistently observed across species, including humans, flies, nematodes, and mice [6]. Patterns of microbial remodeling during aging have been documented in *D. melanogaster*, where older flies typically harbor higher bacterial loads and altered community composition compared to younger individuals [6–8]. The impact of such microbial shifts on lifespan is complex and context-dependent. While some early studies suggested that the absence of microbes shortens lifespan [9], a growing body of evidence indicates that axenic flies often live longer than their conventionally reared counterparts [6,7,10]. These discrepancies likely reflect differences in microbial load, composition, and environmental conditions, including diet. Indeed, microbial abundance has emerged as a stronger determinant of host lifespan than composition alone [6,10]. Transcriptomic analyses further reveal that many host age-associated gene expression changes, such as the downregulation of stress-response genes and upregulation of innate immune genes, are absent in germ-free flies, suggesting that these are adaptive responses to microbial presence during aging [11]. Aging is also associated with structural and functional deterioration of the intestinal epithelium, a key interface between the host and its microbiome. In both flies and mammals, aging leads to increased intestinal permeability, dysregulated immune responses, and hyperproliferation of intestinal stem cells, contributing to dysplasia and loss of barrier integrity [12]. These changes not only compromise nutrient absorption and immune homeostasis but also allow microbial translocation, which can exacerbate systemic inflammation and age-related diseases such as colon cancer and inflammatory bowel disease. In *D. melanogaster*, intestinal barrier dysfunction is a strong predictor of mortality, and flies with leaky guts exhibit significantly reduced lifespans [12]. Maintaining intestinal homeostasis, through balanced intestinal stem cell activity and microbial regulation, is therefore critical for healthy aging. Although *D. melanogaster* does not rely on obligate gut symbionts, the presence of a microbiome has clear fitness consequences. Axenic flies remain viable but exhibit altered metabolism, reduced intestinal aging, and variable effects on lifespan depending on microbial composition and environmental context [13]. Specific bacterial strains influence diverse physiological processes, including insulin signaling, development, and even behavior, underscoring the multifaceted role of the microbiome on host biology [13].

In addition to environmental factors such as diet, host genetics, particularly genes regulating the host immune system, play a significant role in shaping the composition and function of the microbiome [13]. Studies in both humans and model organisms have demonstrated that host genetic variation can influence which microbial taxa colonize the gut, their relative abundances, and their functional interactions with the host [14–16]. For instance, it was demonstrated that in *D. melanogaster* the nutritional impact of the microbiome varies among genetically distinct lines from the *Drosophila* Genetic Reference Panel (DGRP) and host genetic variants were identified and associated with these differences [17]. Findings of highly distinct microbial compositions in *D. melanogaster* lines selected for different stress tolerance traits and longevity also suggest that host and microbiomes may evolve in concert [18]. These findings suggest that host regulatory networks are evolutionarily structured to function in the presence of specific microbial communities, and that dysbiosis can impair host health. Moreover, genetic background of *D. melanogaster* can influence microbial abundance and diversity through mechanisms such as immune modulation, gut physiology, and nutrient availability [13,19]. Consequently, while host genetics clearly modulates microbial interactions, the composition of the microbiome is also subject to considerable environmental and demographic variability. These complexities underscore the need for integrative approaches that consider both genetic and environmental factors when investigating host-microbiome relationships.

*D. melanogaster* is a powerful model for microbiome research due to its genetic tractability, short generation time, and evolutionary conservation of key physiological processes [3,20]. Its gut microbiome is simple, comprising fewer than 30 taxa and it is dominated by the bacterial families *Acetobacteraceae* and *Lactobacillaceae* with consistent representation of species such as *Acetobacter pomorum, A. tropicalis, Lactiplantibacillus plantarum, L. brevis*, and *L. fructivorans* [6,13,17,21–23]. This low-complexity microbiome, shaped by the fly’s diet and acidic gut environment, facilitates reproducible studies of host-microbe interactions and this is a key advantage of using *D. melanogaster* as a model system in microbime research [20,24]. Further, the fly gut shares conserved features in immune signaling, epithelial renewal, and microbial regulation with e.g. humans [4,24,25]. Moreover, the fly gut undergoes age-related changes similar to those in mammals, including dysbiosis and epithelial dysfunction, making it suitable for studying microbiome-driven aging of relevance across taxa [7,12]. Finally, *D. melanogaster* overcomes many limitations of human microbiome studies, such as population heterogeneity, by offering a controlled and reproducible experimental system [2,15]. Together, these features make *D. melanogaster* an ideal model for dissecting the genetic and environmental determinants of microbiome composition and function.

Despite the growing recognition of the microbiome’s role in shaping host phenotypes, the extent to which host genetic variation influences microbiome composition and its interaction with diet remains poorly understood [26]. Individual microbiomes vary widely in both composition and responsiveness to dietary interventions, complicating efforts to generalize their effects on traits related to health and aging. Previous studies have either explored multiple dietary conditions in mice using one or a few genetic lines [27,28], or examined multiple *D. melanogaster* lines in one nutritional environment [13,19,29,30]. However, no study has yet combined both approaches to explore the effects of interactions between genetic variation and diet on the microbiome. This lack of integrative studies leaves a critical gap in our understanding of how genetic variation and diet jointly shape the microbiome and, in turn, influence host aging. Despite this, a growing body of litterature suggest that the microbiome can act as a predictor of late-life performance. For instance, data from the MrOS human cohort, which included community-dwelling older adults aged 78-98 years and followed over a four-year period, revealed that elevated levels of *Bacteroides* were associated with an increased risk of mortality, particularly among individuals aged 85 and older [31]. In contrast, participants with more unique gut microbiome profiles, assessed using Bray-Curtis and weighted UniFrac metrics, exhibited a reduced mortality risk, though this association was only significant in the oldest age group (85+ years). Similarly, it was demonstrated that microbial relative abundances from 21 month old mice could predict healthy aging, defined by a frailty index comprising 31 parameters, for 30 months old mice [32]. In *D. melanogaster*, it has been shown that microbial relative abundances measured at 4 and 25 days of age could predict lifespan, but not other life-history traits such as reproductive output or climbing ability [30]. Although these results are exciting and promising, there is still a lot that we do not know about the importance and predictive power of microbiomes in relation to late life phenotypes. Challenges with humans studies include that they are typically confounded by unctrollable environmental factors and are difficult to replicate [33–35]. Further to that lifespan is also difficult to measure directly in long lived species, so proxy markers like frailty indices and epigenetic clocks are often used [36–38]. Mouse studies typically rely on one or two inbred strains, which simplifies replication but limits the span of genetic diversity assessed and thereby generalizability [27,32]. Moreover, many studies using model organisms focus on either lifespan or healthspan, rarely assessing both or multiple health-related traits [27,35]. Additionally, nearly all published microbiome studies rely on Illumina shortread amplicon sequencing, usually targeting the V3-V4 regions, which offers limited taxonomic resolution and struggles to distinguish closely related microbial strains [27,30–32,35,37]. To overcome some of these these limitations, we used a high number (88) of genetically distinct lines from the *Drosophila* Genetic Reference Panel (DGRP) to investigate how host genetics and diet jointly influence microbiome composition and age-related traits. We investigated the microbiome of young flies exposed to two different diets: a nutritionally rich control diet and a restricted diet. We further investigated lifespan and age-related traits including locomotor activity, dry weight, and heat knockdown time (HKDT), a proxy for heat tolerance, in flies from all lines at different ages. Importantly we designed our experiments in such a way that the host traits locomotor activity, dry weight, and HKDT were assessed on the same individuals. To improve taxonomic resolution, we sequenced the near full 16S rRNA gene. The overarching goal was to obtain a better understanding of whether different genetic backgrounds modulate microbiome responses to dietary interventions and how these interactions affect the aging process. Specifically, we aimed to (i) assess genetic variation in microbiome composition and diversity across dietary environments and (ii) evaluate whether microbiome diversity metrics can serve as predictors of age-related traits in flies under varying dietary conditions.

## Results

We used 88 lines of the DGRP to investigate how host genetic variation and diet shape the gut microbiome and its potential links to age-associated phenotypes. Microbiome profiles were generated from more than 4,700 flies reared on either a nutritionally rich control diet or a restricted diet, and quantitative genetic analyses were applied to estimate broad-sense heritabilities, genetic correlations and genotype-by-diet interactions of microbial features. In parallel, large-scale phenotyping of lifespan, locomotor activity, dry weight, and heat knockdown time (HKDT – a measure of heat tolerance) allowed us to test whether microbiome features could predict variation in aging-related outcomes.

### Impact of Dietary Differences on the Microbiome Composition in Flies

The dataset included 476 microbiome samples, each derived from a pool of ten 6-day-old male flies, with 243 samples from the control diet group and 233 from the restricted diet group, corresponding on average to ∼2.7 replicates per DGRP line per diet. Overall, a significant difference in microbial diversity and composition were observed between flies reared on a control diet and those on a restricted diet (Figure 1). Canonical correspondence analysis (CCA), constrained by diet, revealed that dietary differences explained 0.7% of the variation in CCA1. In contrast, correspondence analysis (CA), representing residual variation not attributed to diet, possible due to line or replicate differences, accounted for 2.4% of the variation in CA1.

**Fig 1.**
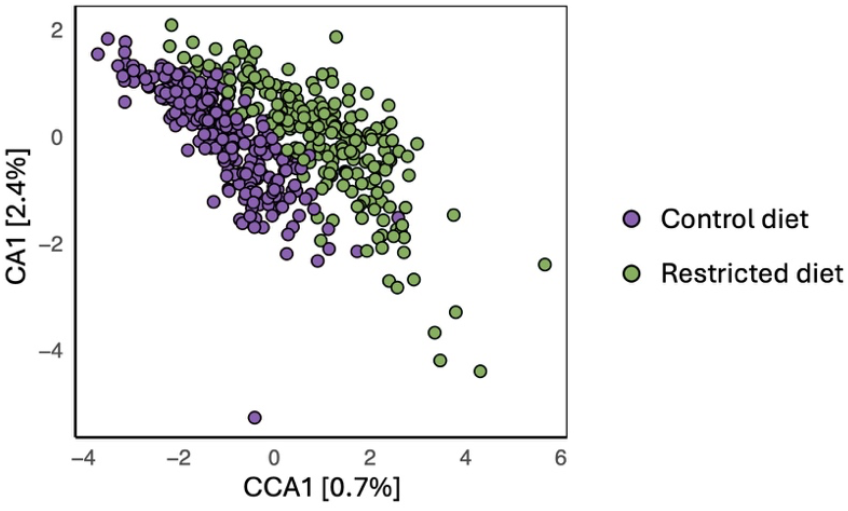
Canonical correspondence analysis and correspondence analysis of microbiome composition. CCA/CA plot illustrating the absolute Operational Taxonomic Units (OTU) abundance between the control (purple) and restricted (green) diets. Euclidean distance was used to assess dissimilarity. Each point represents one microbiome sample (pool of 10 flies).

Principal component analysis (PCA) of the microbiome profiles revealed distinct clustering patterns among fly lines (Figure 2A and 2B). While some lines exhibited clear separation, indicating unique microbial communities, others showed overlapping profiles, suggesting shared microbiome characteristics. Notably, samples from the same line clustered more closely than would be expected by chance, pointing to a consistent within-line microbial signature. Additionally, the degree of variation within each line differed, with some lines displaying tight clustering and others showing broader dispersion, reflecting differences in intra-line microbiome variability and perhaps differences in the ability to regulate the microbiome. The first two principal components explained a greater proportion of variance among lines fed the control diet (34.4%) compared to those on the restricted diet (24.7%).

**Fig 2.**
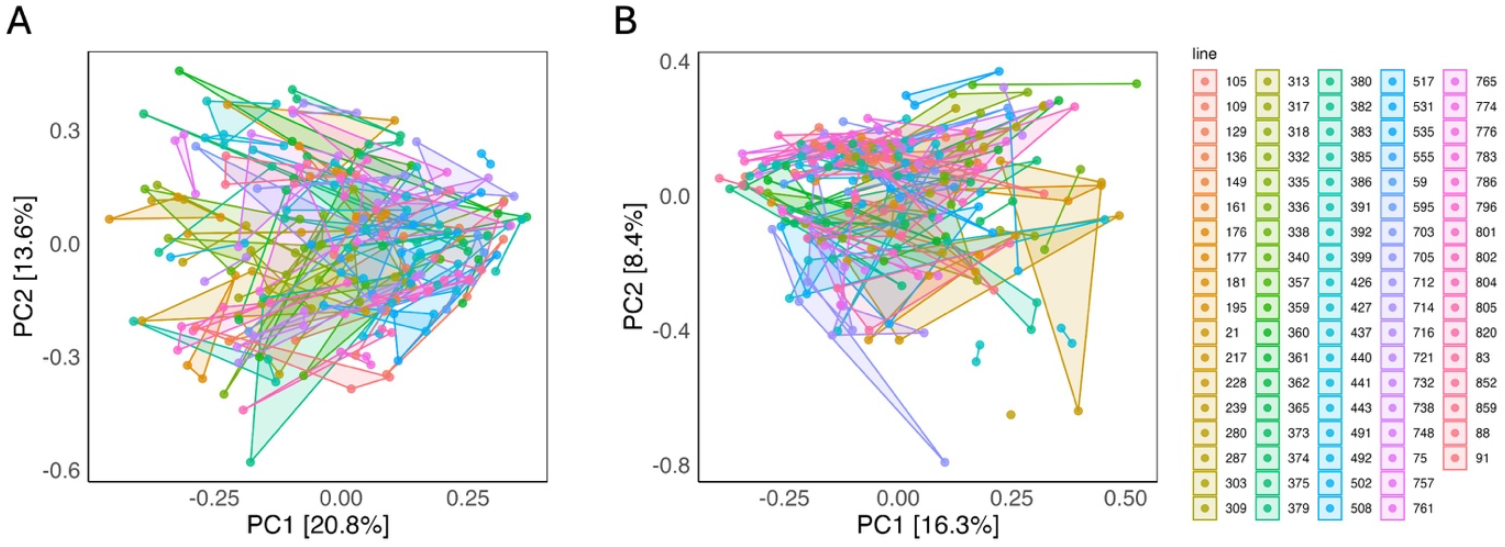
Principal component analysis (PCA) of microbiome composition. PCA based on absolute operational taxonomic units (OTUs) abundance, showing the distribution of samples from DGRP lines under (**A**) the control diet and (**B**) the restricted diet. Distances were calculated using Euclidean metrics. Each point represents one microbiome sample (pool of 10 flies), and points connected by lines correspond to replicate samples from the same DGRP line.

### Host Genetic Background Influences Microbial Community Structure

To assess microbial α-diversity, three metrics were used: number of unique operational taxonomic units (OTUs), Simpson and Shannon indices. A significant difference was observed in the number of unique OTUs, with flies reared on a control diet exhibiting higher richness compared to those on a restricted diet (*p* < 0.0001; Table 1 and S1, and Figure 3A).

**Table 1.** Genetic and environmental contributions to *α*-diversity across diets. Line mean 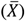, genetic variance 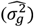, environmental variance 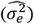 and broad-sense heritabilities 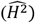 for unique OTUs, Simpson and Shannon indices under control and restricted diet conditions. The table also includes phenotypic (r_p_) and genetic (r_g_) correlations between diets for each α-diversity measure. Values in brackets following the means represent standard errors, while values in brackets following genetic variance, environmental variance, and heritability estimates indicate the confidence intervals.

| Diet | Estimates | Unique OTU | Simpson index | Shannon index |
| --- | --- | --- | --- | --- |
| Control | $\bar{X}$ | 917 [896, 938] | 0.86 [0.858, 0.869] | 3.58 [3.53, 3.63] |
| | $\widehat{\sigma}_g^2$ | 0.12 [0.00, 0.26] | 0.30 [0.16, 0.47] | 0.28 [0.14, 0.45] |
| | $\widehat{\sigma}_e^2$ | 0.72 [0.58, 0.91] | 0.58 [0.47, 0.73] | 0.61 [0.49, 0.77] |
| | $\widehat{H}^2$ | 0.14 [0.01, 0.22] | 0.34 [0.25, 0.40] | 0.31 [0.22, 0.37] |
| Restricted | $\bar{X}$ | 777 [755, 799] | 0.87 [0.866, 0.878] | 3.87 [3.82, 3.93] |
| | $\widehat{\sigma}_g^2$ | 0.06 [0.00, 0.23] | 0.23 [0.06, 0.44] | 0.27 [0.10, 0.48] |
| | $\widehat{\sigma}_e^2$ | 1.00 [0.81, 1.26] | 0.89 [0.71, 1.13] | 0.80 [0.64, 1.01] |
| | $\widehat{H}^2$ | 0.06 [0.00, 0.15] | 0.21 [0.08, 0.28] | 0.25 [0.14, 0.32] |
| | $r_p$ | 0.22 [0.12, 0.32] | 0.46 [0.38, 0.55] | 0.43 [0.34, 0.52] |
| | $r_g$ | 0.95 [0.85, 1.05] | 0.94 [0.85, 1.02] | 0.79 [0.70, 0.88] |

**Fig 3.**
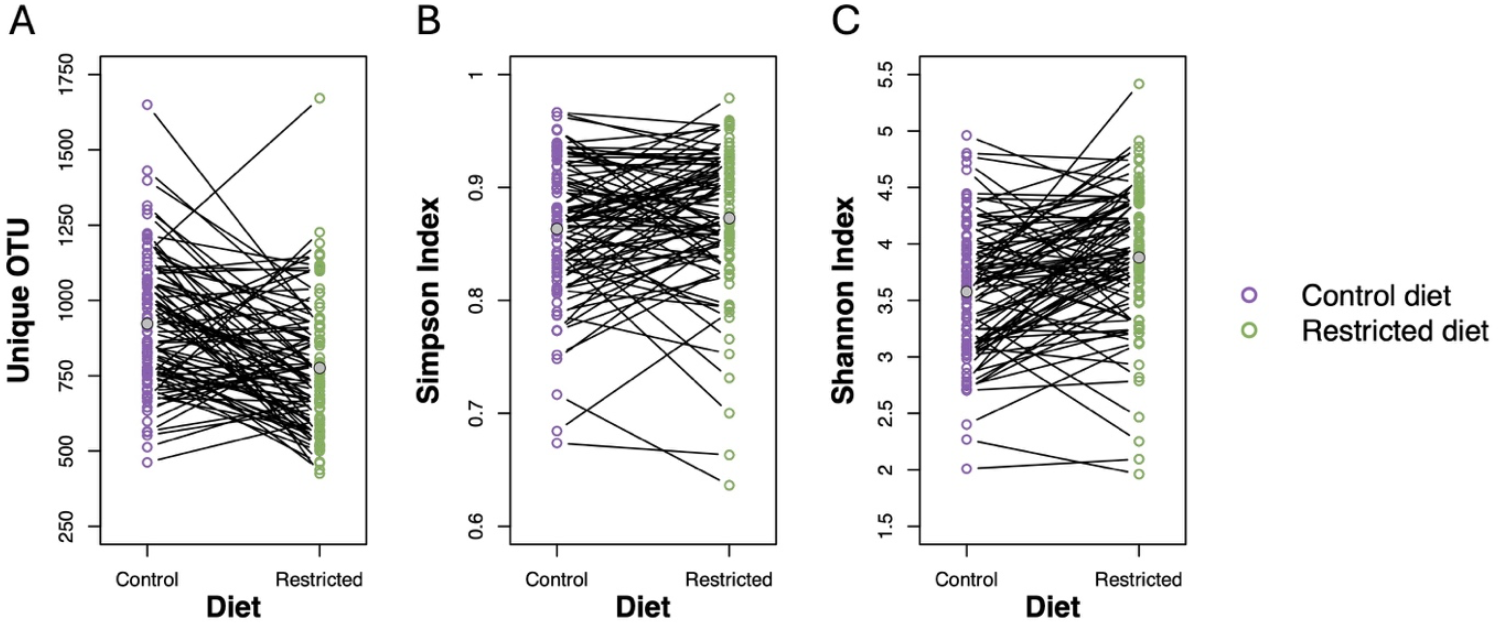
*α*-diversity reaction norm plot. Line means for (**A**) unique OTUs, (**B**) Simpson index, and (**C**) Shannon index in flies fed a control diet (purple) and a restricted diet (green). Each point represents the mean value for an individual DGRP line, with connecting lines indicating the same line across dietary treatments. Grey dots indicate the overall mean across all lines.

In contrast, differences in the Simpson index were only marginal, suggesting broadly similar dominance patterns across dietary groups (*p* = 0.1015, Table 1 and S2, and Figure 3B). However, the Shannon index revealed a significant difference, with flies on the restricted diet displaying a more evenly distributed microbial community, suggesting a more balanced microbiome composition under dietary restriction (*p* = 0.0003; Table 1 and S3, and Figure 3C).

Broad-sense heritability estimates 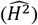 for unique OTUs were 0.14 and 0.06, for the Simpson index 0.34 and 0.21, and for the Shannon index 0.31 and 0.25, under control and restricted diets, respectively (Table 1). The genetic variance for unique OTUs under both dietary conditions, as well as the 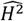 of unique OTUs in flies fed the restricted diet, were not significantly different from zero. In contrast, all other estimates of environmental variance, genetic variance, and 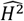 were significantly greater than zero, and no significant differences were observed between diets within each trait (Table 1). Unique OTUs showed a positive phenotypic correlation between diets, while the Simpson and Shannon indices exhibited strong phenotypic correlations between dietary conditions (Table 1). The genetic correlations between control and restricted diets were 0.95 for unique OTUs, 0.94 for the Simpson index, and 0.79 for the Shannon index (Table 1). A significant interaction between diet and DGRP line was observed for both the Simpson (*p* = 0.0418; Table S2) and Shannon (*p* = 0.0248; Table S3) indices, indicating presence of genotype-by-diet interaction (GDI) for these two diversity metrics.

To test whether variation among DGRP lines regulates specific microbes in a diet-dependent manner, we analyzed genera and their corresponding species that exhibited at least 1% relative abundance on at least one diet (Figure 4, S1 and S2, and Table S4 and S5). A clear dominance of *Leuconostoc* was observed across both diets, with particularly high relative abundances (45.83% and 50.91%) in flies on the control diet and restricted diet, respectively. Within this genus *L. pseudomesenteroides* were the most prominent species. *Acetobacter* was the second most abundant genus, with notable species including *A. persici, A. indonesiensis* and *A. cerevisiae*, all showing moderate relative abundances. Other genera such as *Weissella, Lactobacillus* and *Lactiplantibacillus* were present at lower levels, indicating a minor but potentially functionally relevant role. Overall, the ranking of the most abundant genera and species was similar between the control and restricted diets, although their relative abundances differ. For example, *Acetobacter* is more abundant in flies from the the control diet, whereas *Lactiplantibacillus* showed higher abundance in flies from the restricted diet. 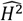 of microbial relative abundance varied across taxa and diets, ranging from 0.21 to 0.57 for genera and 0.18 to 0.74 for species on the control diet, and from 0.25 to 0.54 for for genera and 0.22 to 0.56 for species on the restricted diet (Figure 5 and Table S4 and S5). Phenotypic correlations were moderate overall, spanning 0.06–0.74 for genera and 0.07–0.70 for species. Genetic correlations were consistently higher, ranging from 0.51 to 0.98 for genera and 0.49 to 1.01 for species. All 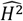 and genetic correlations for both genera and species were non-zero (Figure 5 and Table S4 and S5). Additionally, 87.5% of phenotypic correlations among genera and 90% among species were non-zero. Significant GDI were identified for the bacterial genera *Acetobacter, Lactobacillus* and *Weissella* (*p* = 0.0013, *p* = 0.0310 and *p* = 0.0002, respectively; Table S4) as well as for some species within these genera, including *A. cerevisiae, A. indonesiensis, A. persici, A. sicerae*, and *W. paramesenteroides* (Table S5).

**Fig 4.**
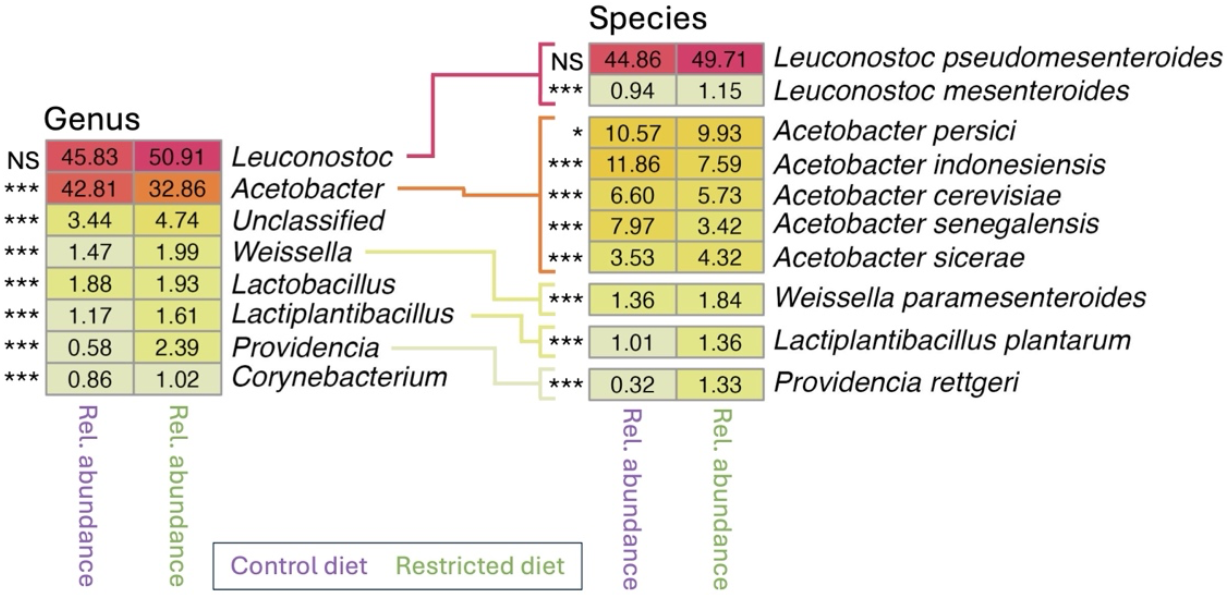
Genera and Species abundance on separate diets. The heatmap on the left displays the genera exhibiting at least 1% relative abundance in flies from at least one of the two dietary conditions, plotted separately for the control (purple) and restricted (green) diets. Species were investigated further if they met three criteria: 1) they belonged to a genus with significantly different abundance between diets, 2) they were known species and 3) they had a relative abundance above 1% on at least one diet. These species are plotted in the heatmap on the right. Relative abundance is represented as a percentage, with dark red indicating the highest relative abundance, yellow indicating medium relative abundance, and olive green indicating the lowest relative abundance. Asterisks (*) denote statistically significant differences in pairwise comparisons between diets, determined using ANOVA (eq. 6), while NS indicates non-significant differences.

**Fig 5.**
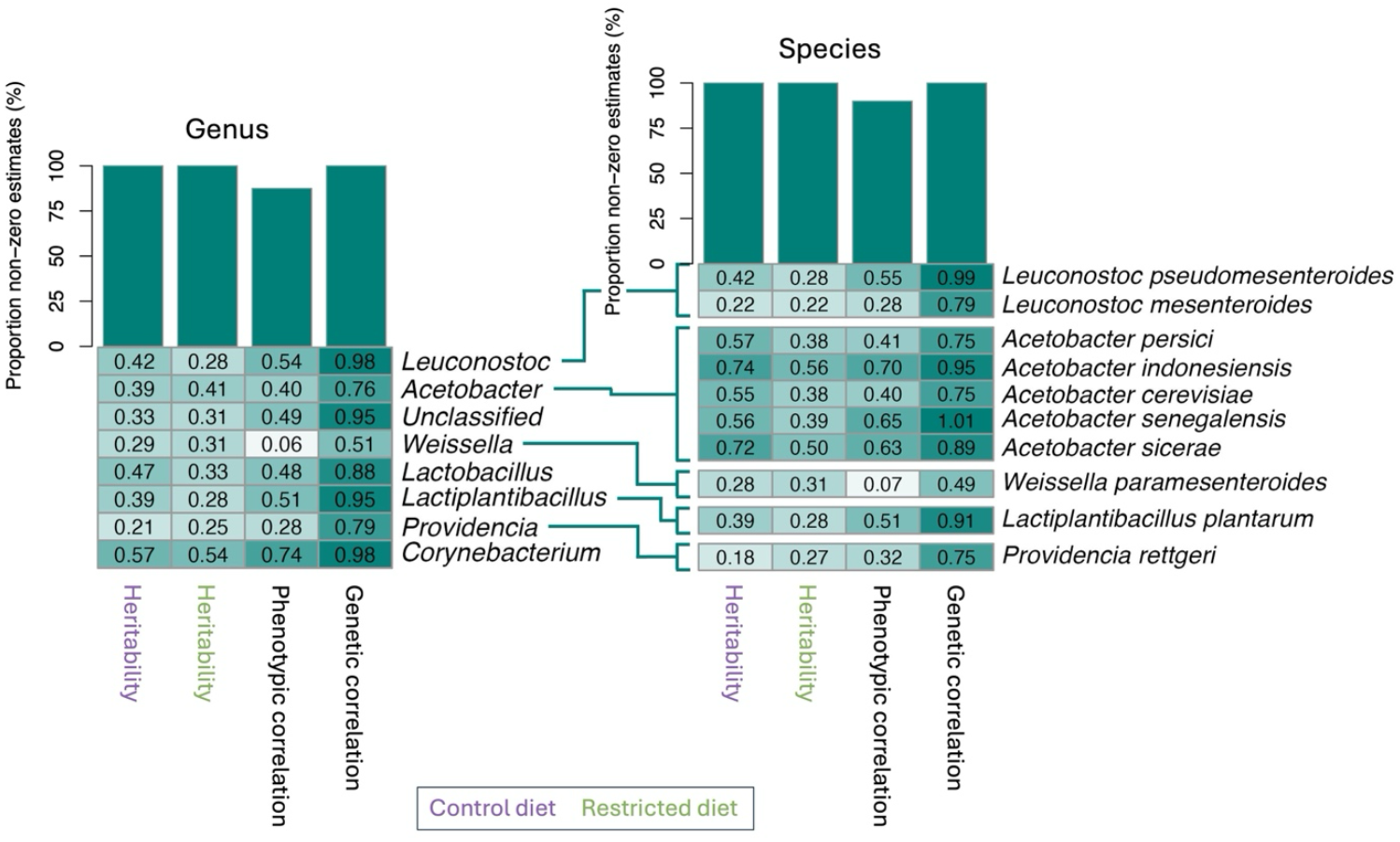
Genetic and environmental contributions to relative abundance of genera and species across diets. The heatmap on the left displays genera with relative abundances above 1% in flies from at least one of the two dietary conditions, plotted separately for the control diet (purple) and restricted diet (green). Species were investigated further if they met three criteria: 1) they belonged to a genus with significantly different abundance between diets, 2) they were known species and 3) they had a relative abundance above 1%. These selected species are displayed in the heatmap to the right. Broad-sense heritability values range from 0 to 1, while phenotypic and genetic correlations range from −1 to 1, represented by a color gradient: dark green indicates a correlation of 1, white indicates zero, and dark red indicates −1. The bar plots above the heatmaps show the proportion of significant non-zero estimates among the genera and species included. None of the broad-sense heritabilities was significantly different between the two diets.

To further explore diet-dependent microbial dynamics, we assessed Spearman correlations of relative abundance for genera with relative abundance above 1% and their respective species across control and restricted diets. We observed a broad spectrum of responses, with several genera and species exhibiting consistently strong positive or negative correlations across both dietary conditions (Figure S3). Notably, the direction of correlations (positive or negative) was largely preserved between diets, with variation primarily observed in the magnitude of the association. For example, *Lactobacillus, Lactiplantibacillus*, and *A. indonesiensis* displayed both positive and negative correlations, whereas *A. cerevisiae* and *A. persici* were predominantly positively correlated, and *Corynebacterium* showed mostly negative correlations. Interestingly, *Leuconostoc* exhibited only positive or neutral correlations on the control diet, but some negative correlations emerged under the restricted diet. These findings suggest that microbial relationships are modulated by dietary context, some genera and species tend to co-occur regardless of diet, while others appear to exclude one another depending on nutritional conditions.

### Lack of Association Between Microbiome Diversity and Age-Related Traits

We next evaluated whether microbiome features based on the samples from the 6 days old male flies could predict age-related traits (lifespan, locomotor activity, heat knockdown time [HKDT], and dry weight). Measurements for locomotor activity, HKDT, and dry weight were taken at 7, 16, 25, and 34 days of age. No statistically significant associations were detected for α-diversity indices (unique OTU, Simpson index or Shannon index; Figure 6, Table S6). Similarly, no significant correlations were found between the relative abundances of genera and species (Table S7–S8). Analyses of ordination scores (PCA1 and PCA2) likewise yielded no significant associations (Table S9).

**Fig 6.**
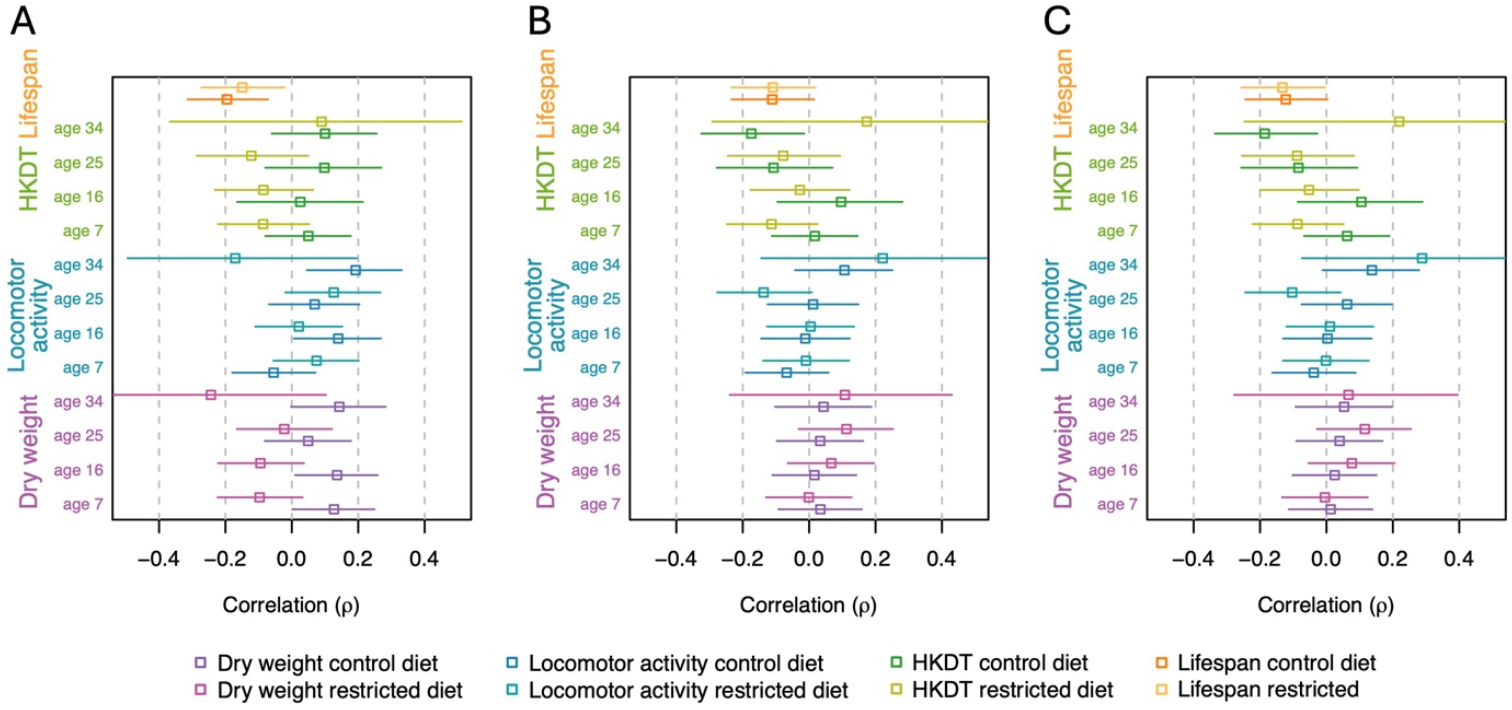
Correlations between *α*-diversities and age-related traits. Age-related traits, dry weight (purple), locomotor activity (blue), heat knockdown time (HKDT; green), and lifespan (orange), were correlated with microbiome-related traits: (A) unique OTUs, (B) Simpson index, and (C) Shannon index. Microbiome data were collected from 6 days old flies, while dry weight, locomotor activity, and HKDT were measured at 7, 16, 25, and 34 days of age. Data are shown for flies fed a control diet (dark shade) and a restricted diet (light shade). Points represent correlation estimates, and error bars indicate corresponding confidence intervals.

## Discussion

The microbiome has highly context-dependent effects on the host, varying with both nutritional environment and genetic background [1–6,14,17,18]. This complexity emphasizes the importance of holistic approaches that integrate both genetic and environmental perspectives when studying host–microbiome relationships. In this study, we examined genetic variation in microbiome composition and diversity across two dietary regimes, and evaluated whether microbiome diversity metrics could serve as predictors of age-related traits in *D. melanogaster*. Using 88 lines from the DGRP tested in both dietary environments, we assessed how host genetic variation and diet influence gut microbiome structure and its potential associations with aging phenotypes. Microbiome data were generated from 476 samples collected from 6 days old flies reared on either a nutrient-rich control diet or a nutrient-poor diet for four days prior to sampling. Simultaneously, we performed large-scale phenotyping of lifespan, locomotor activity, dry weight, and heat knockdown time (HKDT). Importantly, locomotor activity, dry weight, and HKDT were measured on the same individuals and at four time points when flies were 7, 16, 25, and 34 days old, allowing us to investigate the predictive value of microbiome features for age-related traits. This design allowed us to evaluate microbiomes across dietary environments and genetic backgrounds, including impacts of genotype-by-diet interactions on microboimes and host phenotypes.

We employed multiple analytical approaches to assess whether flies fed a nutritionally rich (control) diet harbored a distinctly different microbiome compared to those on a nutritionally poor restricted diet (Figures 1, 3 and 4, and Table 1). The restricted diet used in this study has been shown to reflect a state of malnutrition [39,40], therefore the observed differences in the microbiome between the two diet types may represent a dysbiotic response to nutrient deprivation on the restricted diet rather than an adaptive remodeling. A significant reduction in microbial richness in flies exposed to the restricted compared to the control diet, was revealed by 140 (15.3%) fewer unique OTUs (Figure 3, and Table 1 and S1) and a suppression of *Acetobacter*, including four *Acetobacter* species (Figure 4 and Table S5). This suggest a collapse of the gut microbial ecosystem, with only a subset of resilient taxa persisting. The increased evenness of the microbiome in flies on the restricted diet, as indicated by the 7.5% higher Shannon index (Figure 3, and Table 1 and S3), therefore likely does not reflect a more balanced or stable community, but rather results from a decline in dominant, potentially beneficial taxa and a relative increase in rare, likely non-beneficial taxa. Such patterns are characteristic of malnutrition-induced dysbiosis, which is known to impair gut barrier function, nutrient absorption, and immune homeostasis [6–8,12]. These microbial changes may worsen the negative effects of undernutrition on host health and aging, in contrast to the beneficial effects sometimes associated with controlled dietary restriction. Alternatively, the increase in Shannon index could represent a plastic microbial response aimed at mitigating nutritional stress, potentially through the flourishing of taxa that buffer against malnutrition. Supporting this, *A. persici*, a microbe associated with reduced lifespan via enhanced intestinal stem cell proliferation and immune activation [41], was depleted under the restricted diet. Conversely, *L. plantarum* was enriched (Table S5), a species known to counteract malnutrition and promote longevity by serving as a nutritional source [42]. Nevertheless, the presence of such beneficial microbes may not be sufficient to fully offset the adverse effects of severe nutrient deficiency. In summary, the distinct microbial communities shaped by each diet likely contribute to divergent outcomes in gut health and aging, with the restricted diet fostering either a less favorable microbiome for host resilience or a compensatory microbial shift that remains inadequate to overcome the challenges of malnutrition.

This study explored how microbiome-related traits vary in response to dietary conditions, with a particular focus on the importance of genetic background of the subset of DGRP lines invested here for shaping the microbiome. CCA/CA analyses indicated that diet accounted for only a minor portion of the variation in microbiome profiles, suggesting that other factors, including genetic, play a more dominant role (Figure 1). PCA analysis further revealed that microbiome profiles in flies from the control diet exhibited greater variation between lines, implying that genotype-specific differences in microbiome composition are more pronounced in a nutritionally rich environment (Figure 2). Thus, the control diet may amplify host genetic effects on microbial community structure. This pattern is also reflected in the *α*-diversity measures: flies on the control diet showed relative high 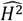 (significantly above zero) for unique OTUs, Simpson, and Shannon indices. Similarly, flies on the restricted diet exhibited 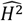 values significantly above zero for the Simpson and Shannon indices. Likewise, 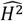 for all tested genera and species were high and significantly greater than zero under both dietary conditions (Figure 5). These consistently high 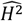 across diet treatments suggest that microbiome-associated traits are under substantial genetic control. This implies that host genomic variation significantly influences microbiome composition and diversity, even under varying environmental conditions. Consequently, the microbiome is not solely shaped by environmental exposure but is also modulated by heritable host factors. While previous studies have also established that both host genetics and environmental factors such as diet influence microbiome composition, the assumption has often been that these influences act in relatively additive or independent ways [43–45]. Our findings challenge this notion by demonstrating that the effect of host genotype on the microbiome is not fixed, but instead varies depending on dietary context. In our study, although many traits exhibited genetic correlations above zero between dietary treatments, the presence of GDI revealed that these correlations were significantly less than unity for several key traits, including Simpson and Shannon indices, and the relative abundances of *Acetobacter, Lactobacillus* and *Weissella*, as well as for species within these genera such as *A. cerevisiae, A. indonesiensis, A. persici, A. sicerae*, and *W. paramesenteroides* (Table S2-S4). This indicates that the genetic control of these traits is not fully conserved across environments. Meaning that the same genetic variant may have different, even opposing, effects on microbiome composition depending on the nutritional environment. For instance, a genotype that promotes the abundance of a beneficial microbial taxon under a nutrient-rich diet may not have the same effect, or may even have a detrimental effect, under dietary restriction. Such context-dependent genetic effects imply that the host–microbiome relationship is governed by a dynamic and conditional architecture, rather than a static one. From a mechanistic perspective, this suggests that different genetic pathways may be activated or suppressed depending on environmental inputs, leading to divergent microbial outcomes. This plasticity complicates efforts to predict or manipulate the microbiome based solely on genetic information, as the environmental context must also be precisely defined and controlled. In practical terms, this has profound implications for microbiome-targeted interventions. For example, in personalized medicine or fecal micobiota transplant therapies, assuming a uniform host genetic influence across dietary conditions could lead to suboptimal or even counterproductive outcomes [26,46]. Similarly, if breeding programs within agriculture aiming to select for favorable microbiome traits, fail to account for GDI this could result in inconsistent trait expression across different feeding regimes or environments [47–51]. Our findings revealing strong GDI are consistent with and extend previous research in mouse models, where significant GDI effects on microbiome traits have been documented [52–54]. In contrast there are only few studies investigating GDI in *D. melanogaster* with microbiome composition as the primary trait. It has been demonstrated that microbial responses can be shaped by GDI [55], while Henry et al. [56] showed through experimental evolution that host genotype can drive shifts in microbiome composition under dietary stress. Nevertheless, the majority of *D. melanogaster* microbiome research tends to examine genetic and environmental influences in isolation or focus on genotype-by-environment interactions in host phenotypes rather than directly assessing microbiome composition [13,14,57]. In summary, the discovery of widespread genotype-by-diet interactions reveals that the genetic regulation of the microbiome is more intricate than previously recognized. It necessitates a shift from static models of host–microbiome interactions to more dynamic frameworks that incorporate environmental modulation of genetic effects. This complexity, while challenging, also opens new avenues for precision microbiome engineering where both host genotype and environmental context are jointly optimized to achieve desired microbial outcomes.

Given that this study demonstrates clear genotype-specific differences in microbiome composition and shows that these differences are heritable, there is a strong rationale for leveraging microbiome data to predict aging-related outcomes. So if heritable microbiome features can be used to predict phenotypes of interest this has perspective in e.g. animal breeding as it suggest that selection for specific early life microbial compositions can be an indirect means to change late life traits of interest. Prior studies have shown that microbiome profiles can predict mortality in humans [31], healthy aging in mice [32], and lifespan in *D. melanogaster* [30], highlighting the use of microbiome data in aging research, even when its predictive power may depend on genetic and environmental context. To evaluate this potential, we investigated correlations between a range of microbiome metrics and age-associated phenotypes. Microbiome features included α-diversity indices (unique OTUs, Simpson, and Shannon), relative abundances of genera and species with 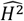 above zero, and ordination scores (PC1 and PC2), all measured in 6 days old flies. Age-related traits assessed were lifespan, locomotor activity, heat knockdown time (HKDT), and dry weight. Notably, locomotor activity, HKDT, and dry weight were measured on the same individuals at four time points; 7, 16, 25, and 34 days old flies, allowing for a robust analysis of microbiome–phenotype associations across developmental stages and dietary conditions. We obtained a clearcut result; none of these correlations were statistical significant, showing that, within this dataset, microbiome-related traits obtained in young flies do not provide a reliable basis for predicting age-related outcomes. Multiple studies have linked gut microbiome profiles to aging such as frailty, survival, and biological age in both humans and mice [32,58], and microbiome transplants from young to old individuals have shown beneficial effects on age-related traits [59–62]. However, evidence that early-life microbiome data can reliably predict long-term aging or lifespan remains limited. For example, Dasari et al. [63], found little to no predictive power of the mortality in baboons and Sultanova et al. [30] found that several life-history traits could not be predicted by the microbiome in *D. melanogaster*. Moreover, other studies have reported no beneficial effects of microbiome transplants on aging traits in *D. melanogaster* [8,64]. This suggests that while the microbiome is shaped by both diet and host genetics, its variation may not directly translate into measurable differences in the investigated age-related traits under the conditions tested here. One possible explanation for the discrepancy between this result and findings in other studies [32,58] is that the genetic variants involved in microbiome regulation may be different from those responsible for maintaining intestinal permeability and gut barrier integrity, such as variants influencing intestinal stem cell proliferation [12]. As a result, some flies may effectively regulate their microbiome, favoring symbionts, while also maintaining a robust gut barrier. Others may similarly regulate their symbionts but have compromised gut barrier function, so even a few pathogenic microbes can reduce their lifespan. A third scenario involves flies unable to regulate their microbiome, resulting in more pathogens, but with a strong gut barrier that mitigates the negative effects on aging. If only the first two scenarios occurred, aging outcomes could be predicted from microbiome composition and gut barrier status. However, the third scenario complicates this relationship, as flies with poor microbiome regulation may still age well if their gut barrier remains intact. Another possible explanation is that microbiome regulation changes with the chronological age of the fly. For instance, a given fly or genotype may regulate its microbiome effectively from a young age and maintain this ability throughout its life. In contrast, another fly may start with optimal regulation but lose this capacity as it ages. A third scenario involves a fly that initially exhibits poor microbiome regulation during early life but either maintains a consistent level of regulation or improves this ability with age, potentially outperforming the second fly in microbiome control during later stages of life. Studies using animal models have demonstrated that the gut microbiome becomes increasingly variable between individuals with age. This variability may reflect a decline in the host’s ability to regulate microbial composition, potentially leading to dysbiosis and associated health deterioration [6–8]. However, fecal microbiota transplantation from young to older individuals has not been shown to reverse aging and may even have harmful effects in *D. melanogaster* [8,64]. This underscores the complexity of microbiome–host interactions and suggests that microbial composition alone is not be sufficient to predict aging outcomes. Our findings align with Wilmanski et al. [65] suggesting that microbial composition alone is a poor predictor of health outcomes, and that microbial function may be a more informative predictor of aging trajectories. In healthy older individuals, microbial metabolites, such as anti-inflammatory compounds, tend to converge, even when microbial taxa diverge. This implies that gut physiology changes with age, and in some cases, the microbiome co-evolves with the host to produce beneficial metabolites. Importantly, individuals with a tightly regulated immune system may maintain control over microbial composition but not necessarily support the adaptive co-evolution of the microbiome required for optimal metabolic function during aging. While for other individuals tight regulation of the immune system is necessary for maintaining optimal metabolic function according to their gut physiology at late age.

Our study shows that genetics strongly influence the microbiome under varying diets, but that microbiome features in young flies does not reliably predict lifespan and age-related traits. These findings are consistent with growing evidence in *D. melanogaster* indicating that fecal microbiota transplants from young individuals do not consistently improve or predict age-related outcomes [8,64]. Therefore, while the microbiome composition remains a promising target in aging research its predictive value seems limited.

## Materials and Methods

### Fly Stock and Nutritional Environments

We utilized 88 DGRP lines, sourced from the Bloomington *Drosophila* Stock Center (NIH P40OD018537, Table S10). The DGRP consists of inbred lines (F=1) derived from a natural population in Raleigh, North Carolina, USA. Each line has undergone 20 generations of full-sib mating, resulting in minimal genetic variation within lines and capturing natural genetic diversity between lines [66,67]. All lines have been genome sequenced at high coverage (∼27X), identifying over 4.5 million genetic variants, including SNPs, indels, and microsatellites.

Prior to experimentation, flies were maintained under standardized conditions: 23 °C, 50% relative humidity, and a 12:12 h light/dark cycle in our laboratory at Aalborg University, Denmark for two generations. They were housed in vials with approximately 10 flies per vial and fed a nutritionally rich control diet composed of dry yeast (60 g/L), sucrose (40 g/L), oatmeal (30 g/L), agar (18 g/L), Nipagen (12 mL/L), and acetic acid (1.2 mL/L) (Table 2).

**Table 2.** Composition of nutritional components in the two diet types. The control diet serves as the reference standard with a defined nutritional value of 100%, while the values for the restrictive diet are presented relative to this baseline.

| Medium types | Yeast | Sucrose | Oat | Agar | Nipagin | Acetic acid | Cellulose |
| --- | --- | --- | --- | --- | --- | --- | --- |
| Control | 60 g/L | 40 g/L | 30 g/L | 18 g/L | 12 mL/L | 1.2 mL/L | 0 g/L |
| Restricted (25%) | 15 g/L | 10 g/L | 7.5 g/L | 18 g/L | 12 mL/L | 1.2 mL/L | 97.5 g/L |

Experimental flies were reared on either this control diet or a restricted diet, created by diluting the control diet to 25% of its nutritional content using indigestible α-cellulose (Product no. 102550125, Sigma-Aldrich, Buchs SG, Switzerland), while keeping agar, Nipagen, and acetic acid concentrations constant across diets (for further details see [39,40]).

### Experimental Design

The age-related trait data used in this study, specifically lifespan, locomotor activity, dry weight, and heat knockdown time (HKDT), were previously analyzed and reported in [40]. In the present study, we build upon that work by incorporating microbiome profiling of flies that were 6 days old.

To generate the F1 generation, approximately 10 adult flies from each DGRP line were transferred to fresh vials every three days, over the course of 5–7 transfers, resulting in 15–20 vials per line. To isolate the effects of adult diet and avoid confounding from developmental nutrition, dietary treatments were applied only during the adult stage. The resulting F1 offspring were pooled into five 250 mL bottles per line, each containing up to 100 flies, to ensure consistent density across lines. These flies were transferred to fresh bottles daily for five consecutive days, yielding up to 25 bottles per line. Upon emergence of the F2 generation, only males were retained for experimentation, while females were discarded. Male sorting continued for four days, and on the fifth day, flies aged 2 days ± 36 hours were allocated into three replicate bottles per diet treatment, each containing 100 flies (or less when this was not possible) (Figure 7). All flies were maintained on the control diet until reaching 2 days of age (± 36 hours), after which they were either continued on the control diet or transferred to the restricted diet.

**Figure 7.**
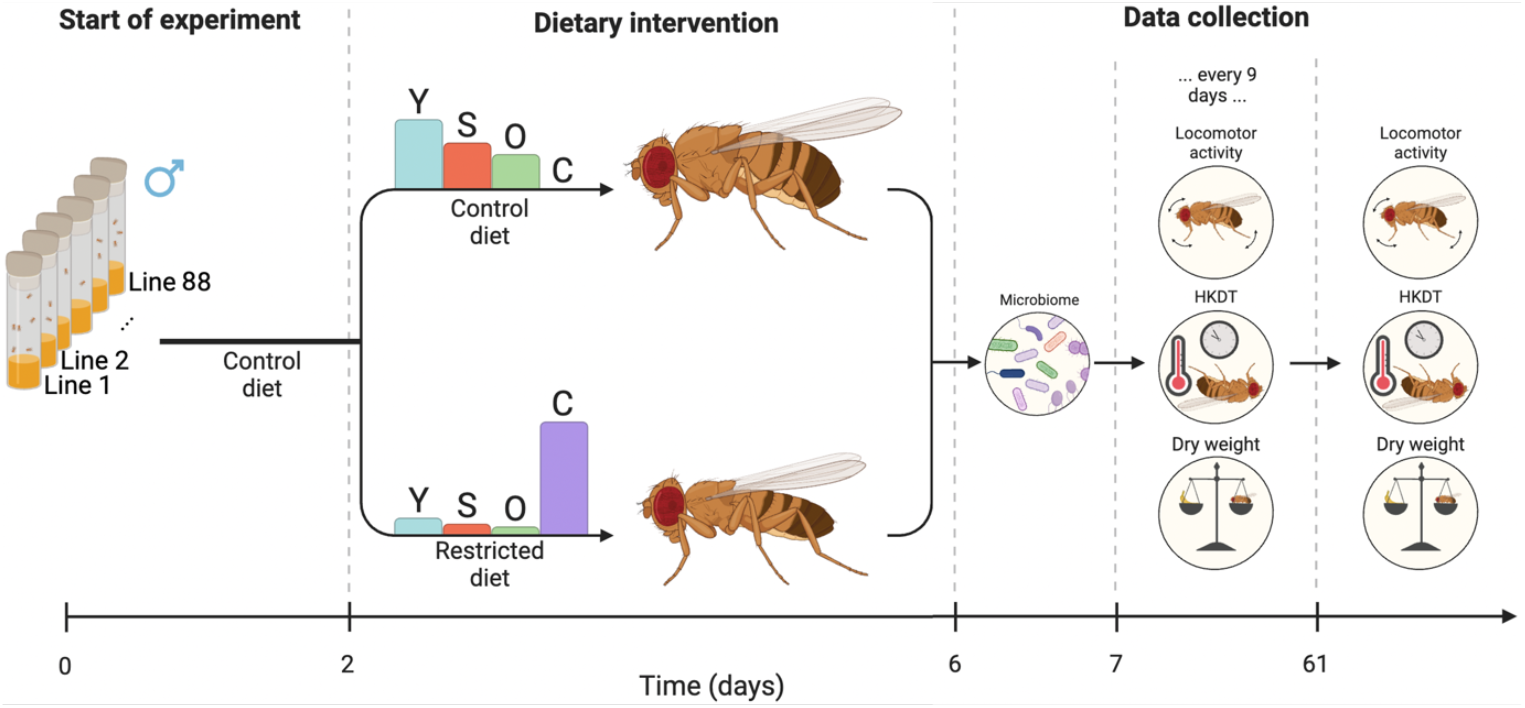
Flowchart of experiment. A total of 88 DGRP lines were maintained at 23°C on a control diet. At 2 days ± 36 hours old, adult flies were transferred to either a control diet (Leeds medium) or a restricted diet. The microbiome was sampled when the flies reached six days of age. Age-related traits, including locomotor activity, HKDT, and dry weight, were assessed at seven days old and then every nine days thereafter. Mortality was recorded every three days, starting from five days old. Bar plots display the relative content of dietary components: yeast (Y), sugar (S), oat (O), and cellulose (C). (Figure is adapted from Bak et al. [2025]).

In total, 48,790 flies were used in the study, with 24,312 reared on the control diet and 24,478 on the restricted diet (Figure 8). Of these, we sequenced the microbiome of 5,040 flies. 4,760 were used for the microbiome analysis after preprocessing of data. Further numerous age-related traits were measured; 6,338 flies were used to measure dry weight, 5,641 for locomotor activity, and 3,666 for HKDT, notably these three phenotypes was measured on the same individual. Importantly, all three age-related traits were measured on the same individuals. These age-related traits were assessed at 7 days of age and then at 9-day intervals until day 61 (N = 1,561 at day 7; N = 1,504 at day 16; N = 1,440 at day 25; N = 1,126 at day 34; N = 530 at day 43; N = 170 at day 52; N = 7 at day 61). Lifespan data were collected from 35,067 flies, 15,904 reared on the control diet (averaging 166 flies per line) and 19,163 on the restricted diet (averaging 196 flies per line). Additionally, 2,345 flies were censored due to escape or due to mortality resulting from physical entrapment, either by becoming embedded in the media or wedged between the flask and the stopper.

**Figure 8.**
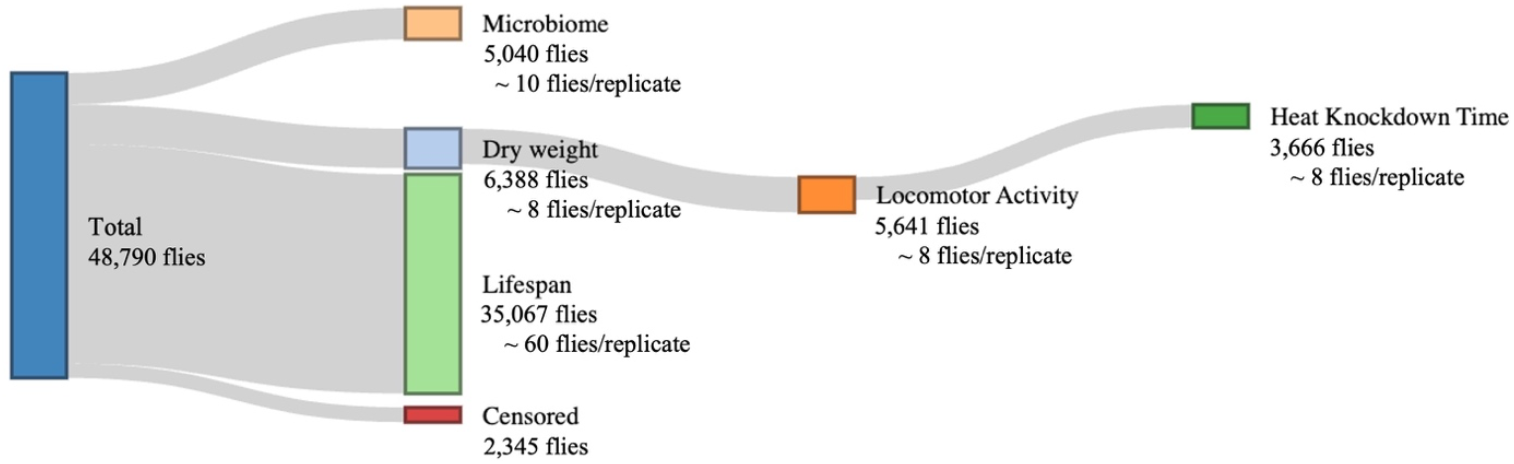
Overview of the total number of flies used across experimental assays. The figure summarizes fly allocation to five main assays: microbiome (5,040 flies; 10 flies per replicate), dry weight (6,388 flies; 8 flies per replicate), locomotor activity (5,641 flies; 8 flies per replicate), heat knockdown time (3,656 flies; 8 flies per replicate), and lifespan (35,067 flies; 60 flies per replicate). Notably, dry weight, locomotor activity, and heat knockdown time were measured on the same individuals. Additionally, 2,945 flies were censored due to escape or entrapment, either by becoming embedded in the medium or lodged between the flask and its stopper.

### Lifespan Assay

For each DGRP line and dietary treatment, we aimed to establish three biological replicates, each consisting of 100 flies housed in individual bottles. While this target was met for most lines, it was not feasible for all. Flies were transferred to fresh bottles containing 60 mL of food medium of either the restricted or or control diet every three days. During each transfer, mortality was recorded, and deceased individuals were removed from the experiment. This process continued until all flies in all replicates had died.

### Locomotor Activity Assay

For the initial locomotor activity assay, eight flies per DGRP line and dietary condition were selected at seven days of age, pooled from all three replicate bottles. These flies were individually placed into 5 mm polycarbonate tubes (TriKinetics Inc., Waltham, MA, USA), each fitted with a water-moistened pipe cleaner at both ends to maintain high humidity. For subsequent time points, we aimed to test 16 flies per line and diet condition. However, due to variation in survival across lines, fewer individuals were available for testing in shorter-lived lines at later stages as reported in [40]. The tubes were inserted into *Drosophila* activity monitors (DAM2, TriKinetics Inc., Waltham, MA, USA), which were positioned in a climate-controlled chamber (Binder KB 400, Binder, Tuttlingen, Germany) maintained at 23 °C. Locomotor activity was recorded as the number of infrared beam crossings per fly, measured every 10 seconds over a 6-hour period (from 9:40 PM to 3:40 AM) (Figure 9). Environmental conditions, including temperature and humidity, were logged every 5 minutes throughout the assay.

**Figure 9.**
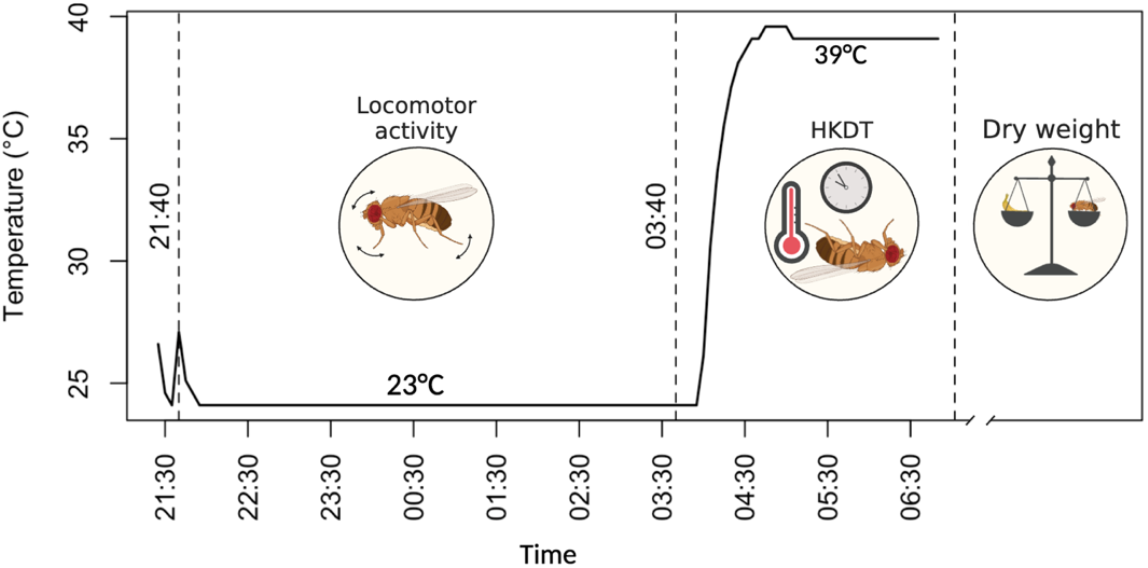
Flowchart of Age-related Traits. All age-related traits were measured on the same individual flies. First locomotor activity was recorded over a six-hour period at 23 °C. This was followed by the heat knockdown time (HKDT) assay, initiated by increasing the temperature to 39 °C, with HKDT defined as the final activity count detected by the *Drosophila* activity monitors. After the assay, flies were stored at −80 °C for subsequent dry weight measurement. The x-axis represents the time of day.

### Heat Stress Tolerance

After completing the locomotor activity assay at 23 °C, flies remained in the *Drosophila* activity monitors and were subsequently exposed to a heat stress condition of 39 °C inside a climate-controlled chamber (Binder KB 400, Binder, Tuttlingen, Germany). Activity was recorded every 10 seconds for an additional 2-hour period (from 3:40 AM to 5:40 AM), during which all flies eventually succumbed to the heat (Figure 9). Heat knockdown time (HKDT) was defined as the time of the last recorded activity count for each individual. Temperature and humidity were continuously monitored at 5-minute intervals throughout the assay. Following the heat exposure, flies were collected and stored individually in Eppendorf tubes at −80 °C for subsequent dry weight measurements.

### Dry Weight

Flies were taken from the freezer and dried in an oven at 60 °C for 48 hours to remove all moisture. Once desiccated, each fly was weighed individually to the nearest 10 µg using a precision balance (Sartorius Quintix35-1S, Sartorius, Göttingen, Germany).

### Sampling, Processing of DNA and Amplicon Sequencing the 16S rRNA Gene

Each flask represented one biological replicate, from which ten flies were sampled at six days of age. When possible, three replicates were collected per line and diet, resulting in a total of 30 flies per line for each diet. Flies were anesthetized using CO_2_, flash-frozen in liquid nitrogen, and subsequently stored at −80 °C. Genomic DNA was extracted from each pooled sample of 10 flies using the DNeasy® Blood & Tissue Kit (Qiagen, Germany), following the manufacturer’s protocol with two modifications: samples were incubated for 2 hours after the addition of Proteinase K and Buffer AL, and DNA was eluted in 80 µL of Buffer AE. DNA concentrations were quantified using the Qubit™ 1× dsDNA HS Assay Kit (Invitrogen, USA) on a Qubit 4 fluorometer (Invitrogen, USA). For bacterial profiling, the 16S rRNA gene was amplified using primers targeting the V1–V8 regions. PCR reactions were performed in duplicate 25 µL volumes, each containing PCRBIO 1X Ultra Mix (PCR BIOSYSTEMS, UK), 400 nM of each primer, 10 ng of template DNA, and nuclease-free water. The thermal cycling conditions included an initial denaturation at 95 °C for 2 minutes; followed by 35 cycles of 95 °C for 15 seconds, 55 °C for 15 seconds, and 72 °C for 90 seconds; and a final extension at 72 °C for 5 minutes. Positive and negative controls were included to verify amplicon quality. PCR products were purified using CleanNGS beads (CleanNA, Netherlands) with a sample-to-bead ratio of 1:0.8, and eluted in 25 µL of nuclease-free water. Library fragment sizes were assessed using the Agilent 4150 TapeStation with D1000/D5000 ScreenTape (Agilent Technologies, USA). Barcoded PCR products were pooled in equimolar amounts, followed by DNA repair, end-prep, adapter ligation, and cleanup. A total of 60 fmol of the final library was loaded onto a MinION R10.4.1 flow cell using the SQK-LSK114 kit with EXP-PBC096, following the manufacturer’s instructions (Oxford Nanopore Technologies, UK). Sequencing was carried out for 72 hours.

### Preparation and Quality Control of Amplicon Sequencing Data

Raw sequencing reads were basecalled and demultiplexed using Dorado (v0.5.0) with the super accuracy model (SUP v4.3.01), applying standard settings in MinKNOW. To ensure precise sample assignment, barcodes were required at both ends of the reads. Further preprocessing was conducted using the ONT-AmpSeq pipeline [68], applying a minimum Q-score threshold of 20 and keeping reads between 1200 and 1600 bp. Operational Taxonomic Units (OTUs) derived from 16S rRNA gene amplification were taxonomically classified using VSEARCH with SINTAX formatted the MiDAS v5.1 reference database [69]. A minimum threshold of 1,000 reads per sample was applied as a cutoff for OTU inclusion, based on rarefaction curves and Good’s coverage [70,71]:

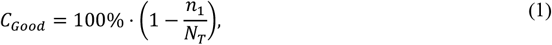

where *n*_*i*_ is the number of singleton OTUs (i.e., OTUs represented by a single sequence), and *N*_*T*_ is the total number of sequences in the sample (Figure S4). To validate the quality of the microbiome sequencing data, the presence of *Wolbachia* was controlled. The known *Wolbachia* infection status of each DGRP line, as reported by [67], was used as a reference. An area under the curve (AUC) analysis was conducted to compare the number of *Wolbachia* sequence reads detected in each sample against the expected infection status (Figure S5). The resulting AUC values were 94.5% for flies maintained on the control diet and 96.1% for those on the restricted diet, indicating high concordance between sequencing data and known *Wolbachia* status. The presence of the endosymbiont *Wolbachia* in insects can influence host fitness and alter the composition or abundance of other microbial taxa, potentially complicating the interpretation of microbiome data [72–75]. To avoid such confounding effects, all sequences identified as *Wolbachia* were excluded prior to conducting the remaining statistical analyses of the microbiome.

### Statistical and Quantitative Genetic Analysis

Rarefaction curves, principal component analysis (PCA) plots and canonical correspondence analysis (CCA/CA) plots were made using the ampvis2 package v. 2.8.9 [76]. To evaluate compositional differences, multiple distance metrics, Chord, Aitchison, Robust Aitchison, Chi-square, and Euclidean, were applied to all PCA and CCA analyses (Figure S6-S8), following recommendations by [77]. As no substantial differences were observed across metrics, Euclidean distance was selected for final visualization. Simpson index, D, was calculated as originally described by [78]:

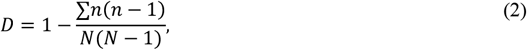

where *n* is the number of individuals of each OTU and *N* is the total number of individuals of all OTU. The Shannon index, H, was formulated by [79]:

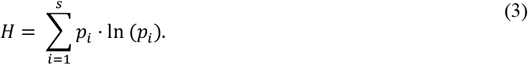

Relative abundance, *R*, was calculated as follows:

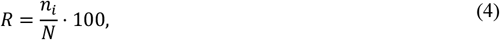

where *n*_*i*_ is the number of reads assigned to genus or species *i*, and *N* is the total number of reads across all genera or species. A rank-based inverse normal transformation (INT) was applied to the unique OTU, Simpson index, Shannon index and relative abundance data to approximate a Gaussian distribution [80,81]. The transformation is defined as:

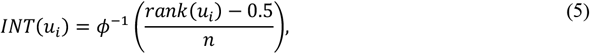

where *Φ*^*-1*^ is the probit (inverse cumulative distribution) function, *rank(u*_*i*_) is the rank of observation *u*_*i*_ and *n* is the total number of samples.

A mixed-model analysis of variance (ANOVA) was conducted using the lme4 package v. 1.1.37 [82] to evaluate how genetic variation among DGRP lines influences differences in unique OTU counts, Simpson index, Shannon index, and the relative abundance of each genus and species within the DGRP population as outlined below:

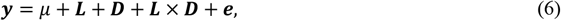

***y*** represents the trait (unique OTUs, Simpson index, Shannon index and relative abundance), *µ* is the fixed intercept corresponding to the overall mean, ***L*** denotes the normally distributed random effect of DGRP line (n = 88), ***D*** is the fixed effect of diet (n = 2), ***L*** × ***D*** is the interaction effect and ***e*** is the normally distributed residual error. A reduced ANOVA model were performed separately for each diet.

Broad-sense heritability, *H*^2^, of the unique OTU, Simpson index, Shannon index and relative abundance was calculated as:

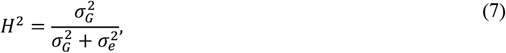

here, 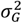 and 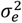 represent the genetic and residual (environmental) variance components, respectively. Standard errors (SE) and confidence intervals (CI) for *H*^2^ were estimated following the approach described in [83]:

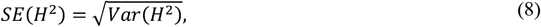

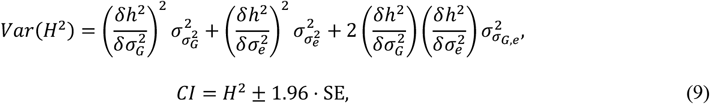

in this context 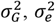, and *σ*_*G e*_ represent elements of the asymptotic covariance matrix. The partial derivatives used in the estimation of *H*^*2*^ are 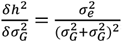, and 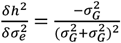. Genetic correlations, *ρ*_*Gij*_, between ages and diets can be estimated following the method described by [84]:

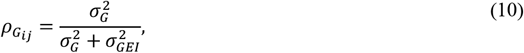

the term 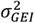 denotes the variance attributed to genotype-by-environment interactions. However, using 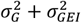 as the denominator when estimating genetic correlations, this approach tends to inflate the denominator due to substantial differences in line variance across environments [84]. As a result, the genetic correlation is underestimated. To correct for this, an alternative formula is used in which the numerator remains unaffected by this bias, providing a more accurate estimate of the genetic correlation:

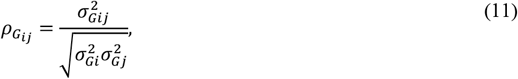

where *i* is the trait on one diet and *j* is the trait on another diet. Spearman correlation coefficients (*ρ*) were used to estimate phenotypic correlations, as described below:

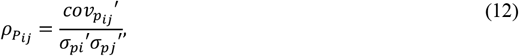

here 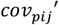 refers to the covariance between the ranked trait *i* measured under a specifc diet and trait *j* measured under a specfic diet while 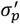 denotes the ranked phenotypic standard deviation. Standard errors and confidence intervals for boh phenotypic and genotypic correlations were estimated following the method described in [85]:

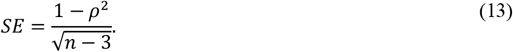

Confidence intervals for the genotypic correlations were likewise estimated following the approach described by [85]:

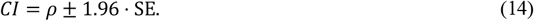

Confidence intervals for the phenotypic correlations were calculated using Fisher’s z-transformation, as described by [86]:

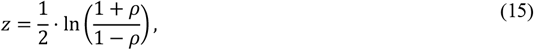

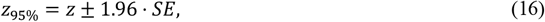

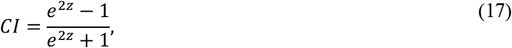

here, z denotes the correlation coefficient using Fisher’s z-transformation.

### Correlation Analysis as a Basis for Trait Prediction

Spearman correlation coefficients were estimated between age-related traits (lifespan, locomotor activity, HKDT, and dry weight) and microbiome-related traits, including *α*-diversity indices (unique OTU, Simpson and Shannon indices), ordination scores (PCA1, and PCA2), and the relative abundance of genera and their respective species with relative abundance above 1%. These correlations were calculated using equation (10), where *cov*_*pij*_′ in this context denotes the covariance between ranked trait *i* of a age-related trait and trait *j* of a microbiome-related trait, both measured under the same dietary condition. The associated standard error, Fisher’s z-transformation and confidence interval were calculated using equation (11), (13-14) and (15), respectively.

## Acknowledgements

We thank Andreas M. Andersen, Anna K. Bak, Christian D. Danielsen, Emilie Battersby, Frederik Kjær, Helle Blendstrup, Jane P. Jakobsen, Jonas B. Wesseltoft, Julie W. Jensen, Madeleine P.S. Madsen, Magnus T. Frantzen, Michael Ørsted, Selma Andersen, Stine F. Laursen, Susan M. Hansen and Timo Kirwa for technical assistance in the laboratory. We further thank Trudy F. C. Mackay and Fabio Morgante for fruitful discussions on the quantitative genetic analyses performed.

## Author contributions

Conceptualization, N.K.B., T.N.K. and P.D.R.; Data Curation, N.K.B. and P.S.S., Formal Analysis, N.K.B., P.D.R. and S.K.Ø.; Funding Acquisition, T.N.K. and P.D.R.; Investigation, N.K.B., T.N.K, S.K.Ø. and P.S.S.; Methodology, N.K.B., T.N.K., P.D.R. and J.L.N.; Project Administration, N.K.B. and T.N.K.; Resources, T.N.K. and P.D.R.; Supervision, T.N.K., and P.D.R.; Visualization, N.K.B.; Writing – Original Draft Preparation, N.K.B., T.N.K. and P.D.R.; Writing – Review and Editing, all authors.

## Conflicts of interest

The authors declare no conflicts of interest.

## Notes

### Competing Interest Statement

The authors have declared no competing interest.

### Funding

- Independent Research Fund Denmark: DFF- 2032-00205A)

