## Supplementary Material 1 for "Host genetics and diet jointly shape the microbiome of *Drosophila melanogaster* but do not predict lifespan or age-related traits"

---

This supplementary file contains:


**Supplementary Figures**

| | |
|---|---|
| Figure S1 | Genera relative abundance for lines on control and restricted diet |
| Figure S2 | Species relative abundance for lines on control and restricted diet |
| Figure S3 | Correlation plots between diets for genera and species |
| Figure S4 | Thredshold for sample cutoff |
| Figure S5 | Receiver Operating Characteristic (ROC) curves comparing *Wolbachia* sequence read counts to expected infection status across dietary treatments |
| Figure S6 | Comparison of distance measures used in CCA |
| Figure S7 | PCA distance measure comparison for flies on a control diet |
| Figure S8 | PCA distance measure comparison for flies on a restricted diet |

**Supplementary Tables**

| | |
|---|---|
| Table S1 | Summary of ANOVA on data of unique OTU |
| Table S2 | Summary of ANOVA on data of Simpson Index |
| Table S3 | Summary of ANOVA on data of Shannon Index |
| Table S4 | Summary of ANOVA on data of relative abundance of genera [**Excel**] |
| Table S5 | Summary of ANOVA on data of relative abundance of species [**Excel**] |
| Table S6 | Correlation between unique OTUs, Simpson Index and Shannon Index with age-related traits [**Excel**] |
| Table S7 | Correlation between relative abundance of genera and age-related traits [**Excel**] |
| Table S8 | Correlation between relative abundance of species and age-related traits [**Excel**] |
| Table S9 | Correlation between PC1 and PC2 scores with age-related traits [**Excel**] |
| Table S10 | Genotype number of the DGRP lines obtained from the Bloomington *Drosophila* Stock Center [**Excel**] |

**A**

| Line | *Leuconostoc* | *Acetobacter* | *Unclassified* | *Weissella* | *Lactobacillus* | *Lactiplantibacillus* | *Providencia* | *Corynebacterium* |
|---|---|---|---|---|---|---|---|---|
| 105 | 44.45 | 39.77 | 2.26 | 0.21 | 9.80 | 0.63 | 0.06 | 0.08 |
| 109 | 58.47 | 29.91 | 8.58 | 1.08 | 0.26 | 0.26 | 0.14 | 0.53 |
| 129 | 44.11 | 49.70 | 2.80 | 1.52 | 0.09 | 0.13 | 0.40 | 0.35 |
| 136 | 54.20 | 35.87 | 3.87 | 2.82 | 0.32 | 0.33 | 0.03 | 1.31 |
| 149 | 53.14 | 33.08 | 2.75 | 0.27 | 0.95 | 0.90 | 6.83 | 0.36 |
| 161 | 42.41 | 33.41 | 2.74 | 11.75 | 5.66 | 0.51 | 0.05 | 0.28 |
| 176 | 52.46 | 34.77 | 3.67 | 0.75 | 3.42 | 1.64 | 0.79 | 0.03 |
| 177 | 7.45 | 74.58 | 3.05 | 0.06 | 9.20 | 2.06 | 0.05 | 0.02 |
| 181 | 12.70 | 73.60 | 2.04 | 7.65 | 1.45 | 1.06 | 0.22 | 0.02 |
| 195 | 48.02 | 48.47 | 1.62 | 0.22 | 0.42 | 0.57 | 0.02 | 0.16 |
| 21 | 59.53 | 37.63 | 1.25 | 0.96 | 0.15 | 0.11 | 0.05 | 0.02 |
| 217 | 32.28 | 55.43 | 2.24 | 0.80 | 0.44 | 0.21 | 7.90 | 0.15 |
| 228 | 31.44 | 59.71 | 2.62 | 1.01 | 1.91 | 1.24 | 0.22 | 0.27 |
| 239 | 3.82 | 87.49 | 1.49 | 0.86 | 3.27 | 0.84 | 0.03 | 0.12 |
| 280 | 25.68 | 68.39 | 2.71 | 0.08 | 0.67 | 1.25 | 0.12 | 0.24 |
| 287 | 21.07 | 45.15 | 25.46 | 0.56 | 0.75 | 0.84 | 0.03 | 4.16 |
| 303 | 50.55 | 23.86 | 5.13 | 3.47 | 0.67 | 0.57 | 0.27 | 13.36 |
| 309 | 24.13 | 61.13 | 4.34 | 2.42 | 2.35 | 2.14 | 0.83 | 0.13 |
| 313 | 61.23 | 21.38 | 4.54 | 2.68 | 1.38 | 1.98 | 0.88 | 2.74 |
| 317 | 42.67 | 46.52 | 3.79 | 2.78 | 1.39 | 1.03 | 0.18 | 0.23 |
| 318 | 34.60 | 53.84 | 3.66 | 2.60 | 1.39 | 1.74 | 0.21 | 0.21 |
| 332 | 30.79 | 59.08 | 4.16 | 0.68 | 1.71 | 1.80 | 0.03 | 0.02 |
| 335 | 40.61 | 44.79 | 11.33 | 0.78 | 0.32 | 0.37 | 0.09 | 0.90 |
| 336 | 31.61 | 60.74 | 2.84 | 1.00 | 1.16 | 1.12 | 0.12 | 0.13 |
| 338 | 22.99 | 52.78 | 4.41 | 14.53 | 1.72 | 1.50 | 0.09 | 0.09 |
| 340 | 58.38 | 29.28 | 4.28 | 2.00 | 1.79 | 0.51 | 0.50 | 0.24 |
| 357 | 37.08 | 49.06 | 8.76 | 0.48 | 0.83 | 0.39 | 0.42 | 0.53 |
| 359 | 59.11 | 33.07 | 2.44 | 2.41 | 0.77 | 0.29 | 0.09 | 0.26 |
| 360 | 66.05 | 27.83 | 2.34 | 0.39 | 0.74 | 0.59 | 0.57 | 0.42 |
| 361 | 70.50 | 23.35 | 2.50 | 0.48 | 0.22 | 0.17 | 0.31 | 0.23 |
| 362 | 52.87 | 30.58 | 2.15 | 0.09 | 5.36 | 2.97 | 0.39 | 3.67 |
| 365 | 28.28 | 59.99 | 2.87 | 3.98 | 1.33 | 1.98 | 0.18 | 0.16 |
| 373 | 43.40 | 47.58 | 2.60 | 0.18 | 1.33 | 0.93 | 0.13 | 0.68 |
| 374 | 45.68 | 49.67 | 2.58 | 0.21 | 0.33 | 0.22 | 0.39 | 0.04 |
| 375 | 43.28 | 50.45 | 2.41 | 1.01 | 0.52 | 0.86 | 0.12 | 0.13 |
| 379 | 59.51 | 33.34 | 2.50 | 0.13 | 0.97 | 1.23 | 0.02 | 0.62 |
| 380 | 19.36 | 69.25 | 3.58 | 1.76 | 1.18 | 1.97 | 0.53 | 0.08 |
| 382 | 25.74 | 68.30 | 2.28 | 1.34 | 0.37 | 0.22 | 0.02 | 0.33 |
| 383 | 48.06 | 40.49 | 3.07 | 2.05 | 1.08 | 1.24 | 0.62 | 0.03 |
| 385 | 13.75 | 72.24 | 3.97 | 3.26 | 2.43 | 1.57 | 0.31 | 0.11 |
| 386 | 70.10 | 16.55 | 4.15 | 2.35 | 0.42 | 0.41 | 0.90 | 1.05 |
| 391 | 83.63 | 13.41 | 1.76 | 0.12 | 0.07 | 0.06 | 0.14 | 0.33 |
| 392 | 58.38 | 31.62 | 2.13 | 0.72 | 0.62 | 0.43 | 0.19 | 5.14 |
| 399 | 27.00 | 58.52 | 3.65 | 0.30 | 3.78 | 3.58 | 0.18 | 0.01 |
| 426 | 62.32 | 34.28 | 1.69 | 0.10 | 0.33 | 0.28 | 0.13 | 0.06 |
| 427 | 39.09 | 56.82 | 2.01 | 0.07 | 0.59 | 0.64 | 0.01 | 0.07 |
| 437 | 43.81 | 52.34 | 1.66 | 0.17 | 0.43 | 0.81 | 0.03 | 0.02 |
| 440 | 24.76 | 36.07 | 2.67 | 0.08 | 19.54 | 11.85 | 0.01 | 0.19 |
| 441 | 66.05 | 23.02 | 2.07 | 6.51 | 0.86 | 0.41 | 0.04 | 0.15 |
| 443 | 40.13 | 43.23 | 2.31 | 0.08 | 0.85 | 1.31 | 0.22 | 3.80 |
| 491 | 44.97 | 50.03 | 2.02 | 0.80 | 0.51 | 0.47 | 0.10 | 0.20 |
| 492 | 15.98 | 76.10 | 1.63 | 1.66 | 2.14 | 1.27 | 0.01 | 0.04 |
| 502 | 44.51 | 33.79 | 4.76 | 0.34 | 2.60 | 1.53 | 7.33 | 2.34 |
| 508 | 85.87 | 8.45 | 2.54 | 0.93 | 0.24 | 0.27 | 0.01 | 0.90 |
| 517 | 61.51 | 27.21 | 4.15 | 0.80 | 1.66 | 2.78 | 0.25 | 0.36 |
| 531 | 60.23 | 17.99 | 5.74 | 0.36 | 1.43 | 0.50 | 0.15 | 1.40 |
| 535 | 47.07 | 33.66 | 10.92 | 1.06 | 2.12 | 2.02 | 0.15 | 1.13 |
| 555 | 52.68 | 37.99 | 4.85 | 0.69 | 0.64 | 0.80 | 0.06 | 0.58 |
| 59 | 66.95 | 22.38 | 3.97 | 1.67 | 0.76 | 0.71 | 0.02 | 2.30 |
| 595 | 40.68 | 49.86 | 3.93 | 0.11 | 1.58 | 1.30 | 0.76 | 0.22 |
| 703 | 46.84 | 42.75 | 3.49 | 1.39 | 1.59 | 0.80 | 0.25 | 0.96 |
| 705 | 41.72 | 47.49 | 3.88 | 2.66 | 0.62 | 1.01 | 0.48 | 0.35 |
| 712 | 76.56 | 16.50 | 2.64 | 0.41 | 0.33 | 0.24 | 0.68 | 1.60 |
| 714 | 46.10 | 51.04 | 1.14 | 0.44 | 0.28 | 0.29 | 0.08 | 0.14 |
| 716 | 34.44 | 50.14 | 6.22 | 0.22 | 2.21 | 1.63 | 0.27 | 1.21 |
| 721 | 13.08 | 57.56 | 13.13 | 0.20 | 2.96 | 1.85 | 0.05 | 7.81 |
| 732 | 78.21 | 16.64 | 2.30 | 0.33 | 0.61 | 0.30 | 0.24 | 0.11 |
| 738 | 18.67 | 59.83 | 8.46 | 0.99 | 0.77 | 1.17 | 0.05 | 1.16 |
| 748 | 16.89 | 65.97 | 8.86 | 0.36 | 0.75 | 0.85 | 0.96 | 1.01 |
| 75 | 60.58 | 28.16 | 4.24 | 0.17 | 1.54 | 1.36 | 0.03 | 1.05 |
| 757 | 12.18 | 74.73 | 3.93 | 1.41 | 1.43 | 2.38 | 0.08 | 0.10 |
| 761 | 62.46 | 23.89 | 5.92 | 0.79 | 0.31 | 0.27 | 0.44 | 3.02 |
| 765 | 35.86 | 49.79 | 3.55 | 3.93 | 2.11 | 1.90 | 0.89 | 0.07 |
| 774 | 53.68 | 41.27 | 1.98 | 0.98 | 0.40 | 0.43 | 0.05 | 0.10 |
| 776 | 35.43 | 41.88 | 4.18 | 16.00 | 0.36 | 0.35 | 0.31 | 0.24 |
| 783 | 65.30 | 27.31 | 4.56 | 0.52 | 0.17 | 0.18 | 0.03 | 0.58 |
| 786 | 51.70 | 41.39 | 2.10 | 1.51 | 0.42 | 0.45 | 1.30 | 0.21 |
| 796 | 30.90 | 52.57 | 3.74 | 0.08 | 8.08 | 0.83 | 0.19 | 0.30 |
| 801 | 51.93 | 42.28 | 3.93 | 0.15 | 0.20 | 0.13 | 0.02 | 0.35 |
| 802 | 8.51 | 78.64 | 4.32 | 0.15 | 2.44 | 1.82 | 0.21 | 0.20 |
| 804 | 30.06 | 41.51 | 5.24 | 0.37 | 8.20 | 1.10 | 0.02 | 10.02 |
| 805 | 59.39 | 31.95 | 4.38 | 1.24 | 0.35 | 0.12 | 0.27 | 0.78 |
| 820 | 54.94 | 38.48 | 2.06 | 0.76 | 0.76 | 1.03 | 1.00 | 0.06 |
| 83 | 37.27 | 50.90 | 1.53 | 0.49 | 0.55 | 0.25 | 0.12 | 0.16 |
| 852 | 78.64 | 17.86 | 1.49 | 0.81 | 0.10 | 0.13 | 0.56 | 0.03 |
| 859 | 34.88 | 58.87 | 2.18 | 0.56 | 0.70 | 0.75 | 0.73 | 0.03 |
| 88 | 46.23 | 48.74 | 1.97 | 0.71 | 0.72 | 0.71 | 0.01 | 0.07 |
| 91 | 31.22 | 62.95 | 1.97 | 0.22 | 0.83 | 1.20 | 0.03 | 0.05 |

**B**

| Line | *Leuconostoc* | *Acetobacter* | *Unclassified* | *Weissella* | *Lactobacillus* | *Lactiplantibacillus* | *Providencia* | *Corynebacterium* |
|---|---|---|---|---|---|---|---|---|
| 105 | 41.28 | 24.51 | 3.82 | 0.37 | 2.40 | 2.04 | 22.96 | 0.19 |
| 109 | 73.06 | 17.56 | 4.72 | 0.20 | 0.83 | 0.46 | 1.15 | 0.44 |
| 129 | 62.67 | 25.03 | 4.44 | 3.21 | 0.50 | 0.23 | 1.75 | 0.74 |
| 136 | 54.62 | 16.08 | 6.57 | 10.36 | 1.55 | 5.34 | 1.93 | 1.45 |
| 149 | 42.08 | 12.20 | 5.72 | 5.43 | 1.12 | 1.68 | 28.26 | 0.38 |
| 161 | 58.81 | 18.55 | 3.67 | 0.10 | 11.15 | 1.10 | 0.43 | 0.37 |
| 176 | 66.59 | 15.76 | 4.22 | 1.10 | 3.29 | 1.64 | 0.75 | 0.04 |
| 177 | 44.16 | 22.25 | 8.91 | 0.85 | 7.03 | 6.62 | 1.00 | 0.72 |
| 181 | 17.01 | 46.44 | 3.76 | 0.22 | 1.19 | 1.68 | 27.44 | 0.04 |
| 195 | 43.28 | 48.61 | 2.82 | 0.68 | 0.32 | 0.43 | 0.08 | 2.77 |
| 21 | 42.50 | 37.62 | 5.60 | 8.58 | 0.41 | 0.26 | 1.89 | 0.25 |
| 217 | 33.46 | 58.33 | 2.64 | 0.57 | 1.58 | 1.05 | 0.38 | 0.15 |
| 228 | 9.63 | 75.12 | 5.03 | 2.18 | 1.80 | 1.83 | 0.87 | 0.21 |
| 239 | 7.52 | 85.10 | 2.45 | 0.08 | 1.19 | 0.96 | 0.02 | 0.21 |
| 280 | 52.21 | 36.00 | 5.38 | 1.76 | 0.83 | 1.11 | 0.22 | 0.30 |
| 287 | 14.79 | 44.25 | 28.23 | 0.63 | 0.97 | 1.32 | 1.14 | 5.29 |
| 303 | 47.01 | 43.65 | 2.88 | 0.37 | 0.43 | 0.63 | 0.75 | 3.53 |
| 309 | 27.40 | 33.76 | 4.90 | 3.67 | 15.93 | 10.03 | 0.09 | 0.71 |
| 313 | 58.97 | 24.64 | 4.70 | 1.97 | 1.73 | 1.63 | 0.16 | 2.45 |
| 317 | 71.52 | 23.29 | 2.11 | 0.38 | 0.59 | 0.86 | 0.07 | 0.30 |
| 318 | 28.79 | 65.33 | 2.79 | 0.37 | 0.82 | 0.65 | 0.18 | 0.10 |
| 332 | 37.52 | 54.37 | 2.25 | 0.21 | 2.02 | 1.89 | 0.03 | 0.03 |
| 335 | 25.46 | 25.52 | 25.70 | 1.59 | 5.73 | 4.42 | 1.83 | 3.05 |
| 336 | 19.70 | 75.17 | 2.58 | 0.16 | 0.66 | 0.65 | 0.28 | 0.01 |
| 338 | 69.84 | 17.71 | 5.74 | 0.67 | 1.05 | 1.23 | 0.46 | 0.77 |
| 340 | 35.22 | 40.71 | 12.17 | 1.37 | 2.76 | 2.04 | 0.20 | 0.61 |
| 357 | 22.53 | 68.99 | 3.59 | 0.34 | 1.35 | 0.67 | 0.06 | 0.31 |
| 359 | 39.75 | 49.52 | 5.05 | 1.25 | 0.61 | 0.55 | 0.91 | 0.91 |
| 360 | 52.46 | 33.83 | 4.94 | 1.68 | 0.93 | 0.80 | 1.40 | 1.67 |
| 361 | 38.59 | 56.09 | 2.14 | 0.21 | 0.49 | 0.58 | 0.98 | 0.11 |
| 362 | 63.35 | 13.02 | 3.14 | 7.11 | 0.45 | 0.45 | 1.52 | 9.80 |
| 365 | 66.91 | 13.83 | 7.24 | 7.16 | 0.64 | 0.78 | 1.20 | 0.92 |
| 373 | 22.25 | 46.66 | 12.56 | 5.55 | 1.43 | 1.83 | 1.77 | 3.54 |
| 374 | 45.43 | 12.43 | 7.06 | 27.38 | 1.42 | 1.03 | 0.62 | 0.96 |
| 375 | 69.96 | 17.18 | 5.09 | 0.58 | 1.40 | 2.55 | 1.12 | 0.33 |
| 379 | 71.11 | 23.46 | 2.18 | 1.40 | 0.33 | 0.29 | 0.35 | 0.23 |
| 380 | 60.58 | 12.79 | 7.32 | 3.77 | 2.08 | 2.94 | 2.23 | 0.97 |
| 382 | 62.98 | 27.81 | 4.83 | 1.13 | 0.31 | 0.48 | 0.04 | 0.71 |
| 383 | 18.38 | 58.91 | 6.59 | 2.47 | 3.33 | 4.37 | 0.78 | 0.12 |
| 385 | 48.12 | 41.97 | 3.71 | 1.96 | 0.86 | 1.22 | 0.42 | 0.07 |
| 386 | 72.22 | 21.71 | 3.19 | 0.31 | 0.30 | 0.35 | 0.26 | 0.94 |
| 391 | 86.59 | 10.48 | 1.18 | 0.29 | 0.37 | 0.29 | 0.16 | 0.16 |
| 392 | 64.25 | 14.59 | 6.04 | 5.29 | 1.51 | 0.80 | 2.01 | 2.41 |
| 399 | 19.14 | 62.13 | 5.21 | 0.29 | 3.67 | 3.46 | 1.46 | 0.41 |
| 426 | 80.17 | 11.73 | 3.50 | 0.46 | 1.18 | 1.23 | 0.03 | 0.12 |
| 427 | 50.21 | 39.65 | 3.68 | 2.01 | 1.04 | 1.21 | 0.45 | 0.03 |
| 437 | 63.77 | 31.10 | 2.27 | 0.28 | 0.64 | 0.91 | 0.23 | 0.12 |
| 440 | 9.50 | 74.09 | 6.19 | 1.89 | 1.84 | 2.85 | 0.12 | 0.26 |
| 441 | 69.63 | 15.44 | 5.10 | 3.71 | 1.82 | 1.48 | 0.42 | 0.12 |
| 443 | 55.47 | 25.67 | 5.91 | 1.91 | 1.29 | 1.70 | 0.02 | 6.03 |
| 491 | 50.59 | 22.83 | 8.91 | 3.65 | 2.74 | 2.31 | 1.98 | 1.09 |
| 492 | 28.77 | 50.31 | 7.43 | 2.99 | 2.99 | 3.33 | 1.26 | 0.11 |
| 502 | 15.36 | 60.81 | 6.09 | 4.85 | 2.13 | 1.71 | 3.23 | 1.08 |
| 508 | 71.84 | 21.99 | 2.56 | 1.35 | 0.22 | 0.21 | 0.36 | 0.49 |
| 517 | 49.60 | 43.05 | 2.50 | 1.10 | 0.76 | 0.95 | 0.13 | 0.09 |
| 531 | 45.15 | 39.53 | 6.61 | 0.75 | 0.94 | 0.85 | 0.09 | 2.03 |
| 535 | 36.83 | 34.56 | 11.51 | 4.34 | 2.57 | 2.20 | 2.52 | 1.64 |
| 555 | 39.67 | 42.27 | 7.73 | 0.70 | 2.09 | 2.36 | 0.96 | 0.81 |
| 59 | 42.90 | 40.11 | 5.30 | 3.48 | 2.20 | 2.26 | 0.05 | 0.81 |
| 595 | 49.10 | 22.28 | 9.02 | 2.49 | 2.80 | 4.51 | 3.02 | 1.05 |
| 703 | 58.61 | 24.66 | 5.83 | 0.73 | 2.47 | 1.05 | 0.07 | 3.44 |
| 705 | 61.96 | 32.07 | 2.27 | 1.24 | 0.25 | 0.29 | 0.54 | 0.31 |
| 712 | 64.52 | 22.22 | 4.61 | 2.68 | 0.35 | 0.31 | 1.30 | 1.33 |
| 714 | 57.46 | 35.75 | 2.36 | 1.30 | 0.80 | 0.91 | 0.29 | 0.13 |
| 716 | 65.98 | 16.37 | 6.89 | 0.85 | 1.68 | 0.61 | 2.68 | 1.68 |
| 721 | 14.41 | 34.94 | 7.59 | 3.21 | 14.27 | 21.09 | 0.13 | 0.67 |
| 732 | 55.03 | 20.05 | 4.85 | 2.22 | 2.23 | 1.67 | 1.00 | 1.30 |
| 738 | 24.55 | 53.03 | 9.44 | 1.62 | 1.19 | 1.37 | 0.58 | 2.46 |
| 748 | 64.05 | 17.99 | 13.46 | 0.23 | 0.31 | 0.32 | 1.05 | 1.19 |
| 75 | 76.01 | 5.08 | 7.50 | 2.48 | 2.23 | 1.35 | 1.40 | 0.82 |
| 757 | 47.70 | 27.61 | 7.30 | 7.32 | 1.18 | 1.47 | 0.11 | 0.94 |
| 761 | 57.10 | 13.67 | 9.09 | 2.67 | 0.87 | 0.63 | 2.08 | 10.10 |
| 765 | 54.43 | 22.25 | 4.19 | 1.55 | 5.59 | 7.92 | 0.22 | 0.29 |
| 774 | 52.56 | 37.36 | 3.20 | 0.72 | 1.92 | 1.59 | 0.32 | 0.17 |
| 776 | 48.69 | 31.63 | 6.53 | 7.86 | 1.61 | 0.93 | 0.07 | 0.18 |
| 783 | 76.22 | 11.48 | 6.31 | 0.84 | 0.78 | 0.67 | 0.75 | 0.94 |
| 786 | 65.56 | 18.19 | 6.93 | 0.85 | 0.39 | 0.21 | 1.98 | 1.24 |
| 796 | 60.43 | 19.43 | 8.11 | 5.04 | 0.93 | 0.83 | 1.19 | 0.64 |
| 801 | 66.61 | 6.47 | 10.65 | 6.48 | 2.21 | 2.59 | 0.31 | 1.14 |
| 802 | 32.73 | 36.90 | 5.39 | 12.20 | 3.37 | 3.49 | 1.19 | 0.23 |
| 804 | 70.11 | 13.56 | 5.54 | 0.28 | 0.90 | 0.92 | 1.52 | 4.56 |
| 805 | 53.02 | 16.14 | 15.81 | 0.51 | 0.55 | 0.45 | 7.09 | 1.41 |
| 820 | 30.24 | 21.15 | 4.79 | 0.41 | 2.98 | 3.60 | 34.36 | 0.06 |
| 83 | 41.20 | 36.62 | 5.75 | 4.20 | 4.23 | 1.58 | 0.05 | 0.29 |
| 852 | 89.90 | 5.26 | 2.57 | 0.22 | 0.18 | 0.18 | 0.40 | 0.22 |
| 859 | 56.65 | 36.12 | 3.25 | 0.35 | 0.51 | 0.35 | 1.06 | 0.08 |
| 88 | 48.98 | 32.24 | 3.94 | 0.83 | 6.31 | 2.91 | 0.17 | 0.10 |
| 91 | 42.10 | 18.03 | 6.04 | 18.17 | 3.17 | 5.97 | 1.47 | 0.27 |

**Fig S1. Genera relative abundance for lines on control and restricted diet.** The heatmaps shows genera for the separate DGRP lines on (**A**) the control diet and (**B**) the restricted diet. Genera was kept if they exhibited at least 1% relative abundance in flies from at least one of the two dietary conditions. Relative abundance is represented as a percentage, with dark red indicating the highest abundance, yellow indicating medium abundance, and olive green indicating the lowest abundance.

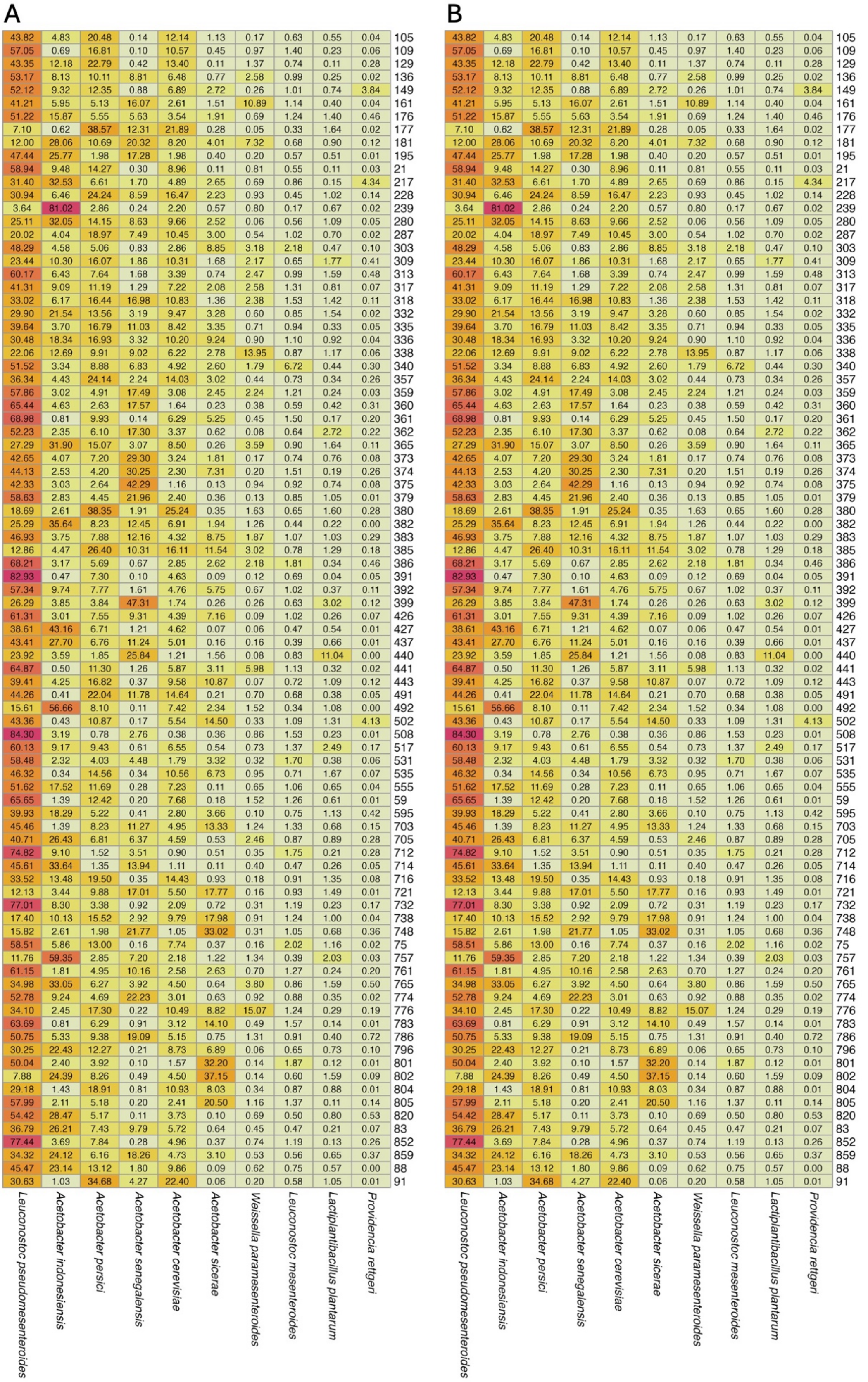


**Fig S2. Species relative abundance for lines on control and restricted diet.** The heatmaps shows species for the separate DGRP lines on (**A**) the control diet and (**B**) the restricted diet. Species was kept if they exhibited at least 1% relative abundance in flies from at least one of the two dietary conditions. Relative abundance is represented as a percentage, with dark red indicating the highest abundance, yellow indicating medium abundance, and olive green indicating the lowest abundance.

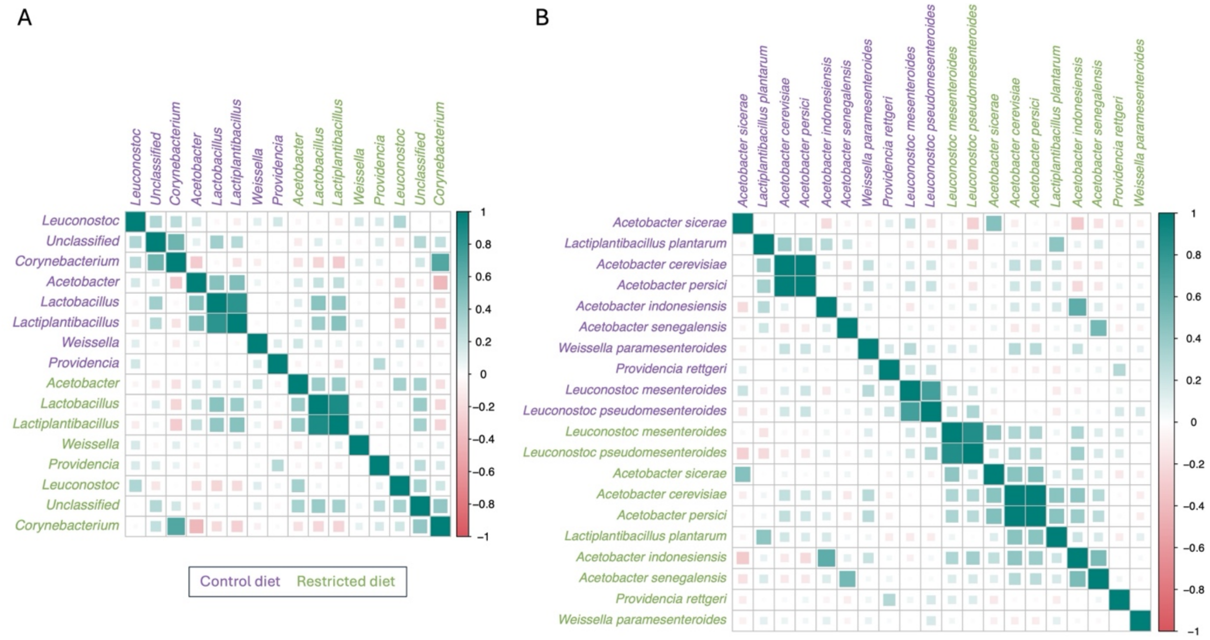


**Fig S3. Correlation plots between diets of genera and species.** Spearman correlations coefficient for the relative abundance of (A) genera and (B) species, with relative abundances above 1%, between control diet (purple) and restricted diet (green). The color and size of the markers represent the strength and direction of the correlations, green indicates positive correlations, red indicates negative ones, and larger markers denote stronger correlations.

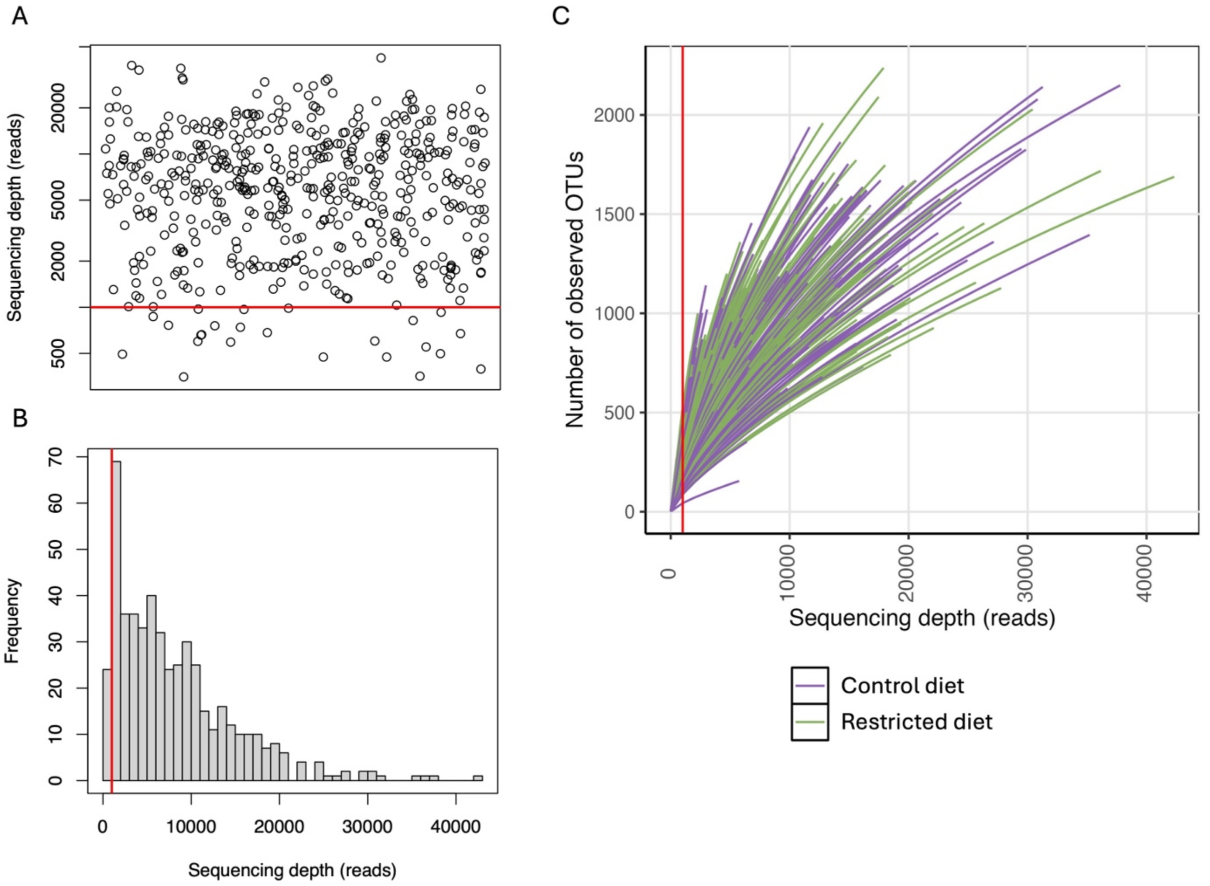


**Fig S4. Thredshold for sample cutoff.** Sequencing depth was vizualised using (**A**) a jitter plot, (**B**) a histogram and (**C**) a rarefaction curve for flies maintained on the control diet (purple) and on the restricted diet (green). Red line indicates a sequencing depth of 1000 reads.

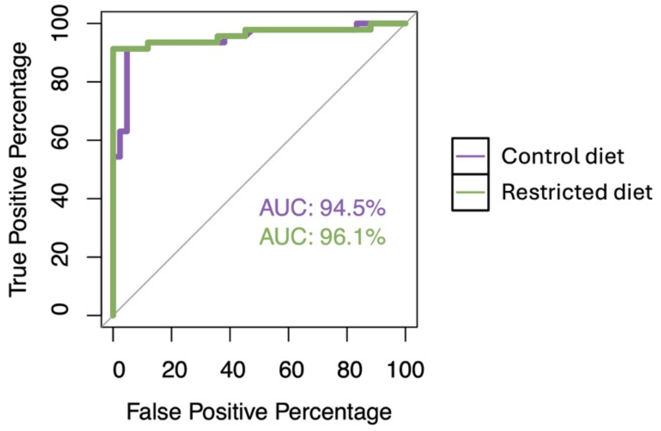


**Fig S5. Receiver Operating Characteristic (ROC) curves comparing *Wolbachia* sequence read counts to expected infection status across dietary treatments.** The ROC curve for flies maintained on the control diet (purple) yield an Area Under the Curve (AUC) of 94.5%, while the curve for flies on the restricted diet (green) with an AUC of 96.1%.

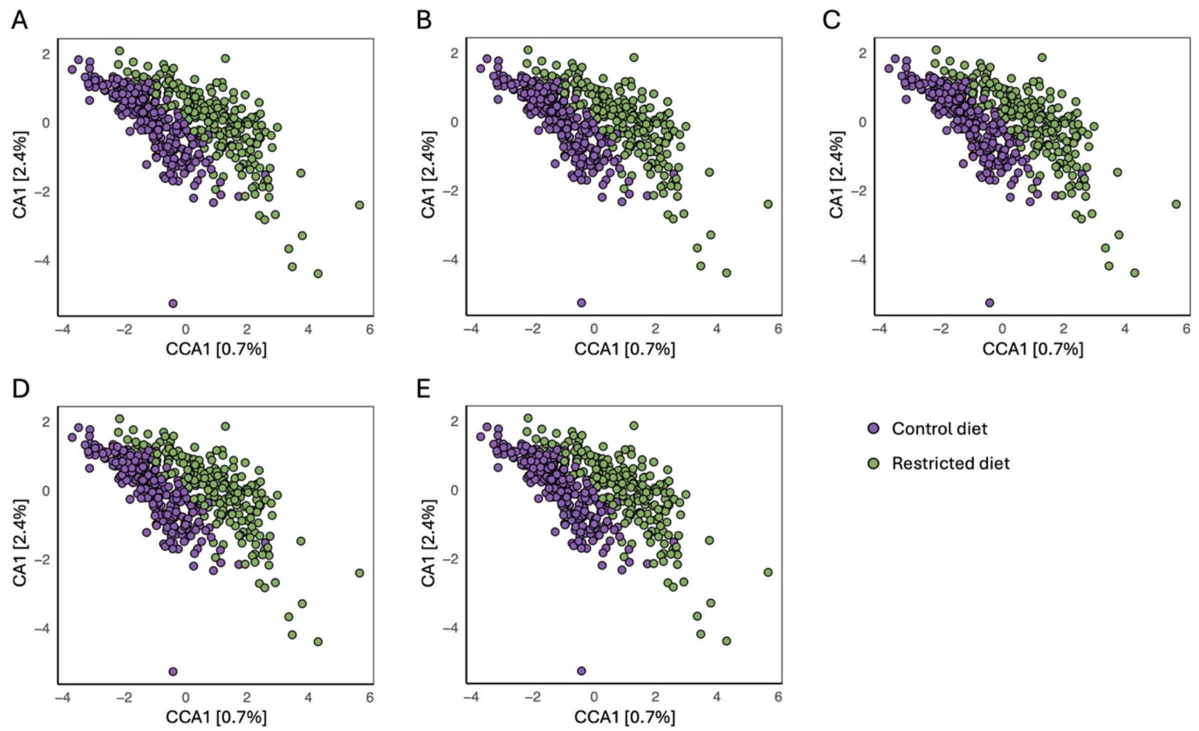


**Fig S6. Comparison of distance measures used in CCA/CA plot.** Panels show CCA/CA plots based on five different distance metrics: (A) Aitchison, (B) Chi-square, (C) Chord, (D) Euclidean, and (E) Robust Aitchison. Each plot displays the first canonical axis (CCA1) on the x-axis and the first correspondence axis (CA1) on the y-axis, with the percentage of variance explained indicated in brackets. Data points are color-coded by diet group: control (purple) and restricted (green).

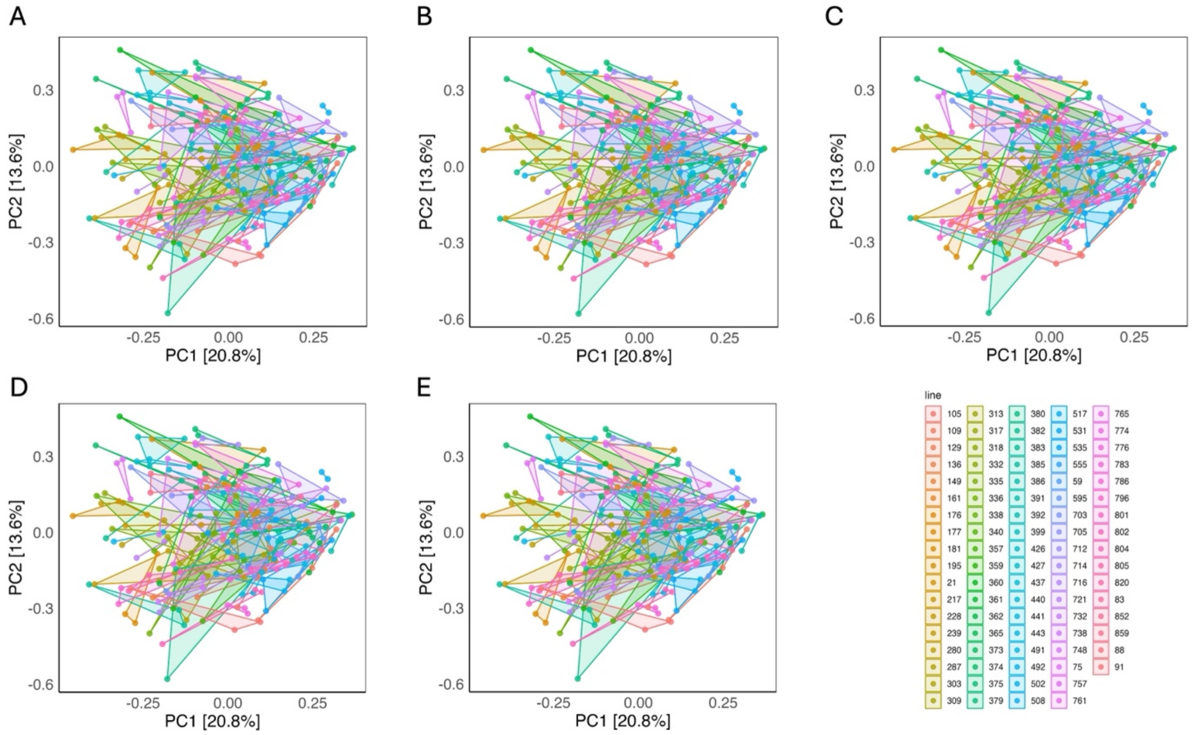


**Fig S7. PCA distance measure comparison for flies on a control diet.** PCA plots are shown for five different distance metrics: (A) Aitchison, (B) Chi-square, (C) Chord, (D) Euclidean, and (E) Robust Aitchison. Each plot displays the first two principal components (PC1 and PC2), with the percentage of variance explained indicated on each axis. The color gradient represent DGRP lines.

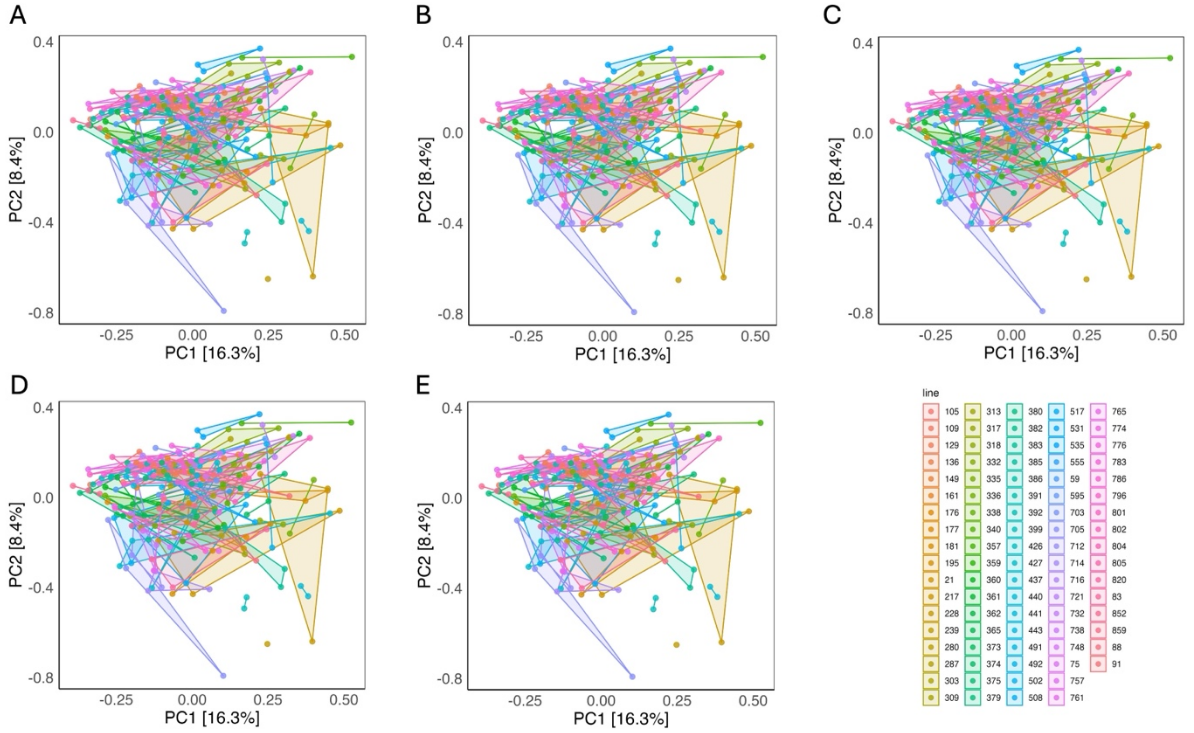


**Fig S8. PCA distance measure comparison for flies on a restricted diet.** PCA plots are shown for five different distance metrics: (A) Aitchison, (B) Chi-square, (C) Chord, (D) Euclidean, and (E) Robust Aitchison. Each plot displays the first two principal components (PC1 and PC2), with the percentage of variance explained indicated on each axis. The color gradient represent DGRP lines.

**Table S1.** Overview of full and reduced ANOVA models evaluating unique OTUs in DGRP flies under control and restricted dietary conditions. Diet is treated as a fixed effect, while DGRP line is modeled as a random effect.

| Condition | Source | DF | Mean Sq | F Value | *p*-value |
|---|---|---|---|---|---|
| Control diet | L | 1 | 3.90 | 4.68 | 0.0315 |
| | $\epsilon$ | 241 | 0.83 | - | - |
| Restricted diet | L | 1 | 8.16 | 7.90 | 0.0054 |
| | $\epsilon$ | 231 | 1.03 | - | - |
| Both diets | L | 1 | 11.67 | 12.54 | 0.0004 |
| | D | 1 | 23.33 | 25.07 | <0.0001 |
| | L×D | 1 | 0.39 | 0.420 | 0.5172 |
| | $\epsilon$ | 472 | 0.93 | - | - |

**Table S2.** Overview of full and reduced ANOVA models evaluating Simpson Index in DGRP flies under control and restricted dietary conditions. Diet is treated as a fixed effect, while DGRP line is modeled as a random effect.

| Condition | Source | DF | Mean Sq | F Value | *p*-value |
|---|---|---|---|---|---|
| Control diet | L | 1 | 1.71 | 1.96 | 0.1629 |
| | $\epsilon$ | 241 | 0.87 | - | - |
| Restricted diet | L | 1 | 2.45 | 2.20 | 0.1394 |
| | $\epsilon$ | 231 | 1.11 | - | - |
| Both diets | L | 1 | 0.03 | 0.03 | 0.8552 |
| | D | 1 | 2.67 | 2.69 | 0.1015 |
| | L×D | 1 | 4.13 | 4.16 | 0.0418 |
| | $\epsilon$ | 472 | 0.99 | - | - |

**Table S3.** Overview of full and reduced ANOVA models evaluating Shannon Index in DGRP flies under control and restricted dietary conditions. Diet is treated as a fixed effect, while DGRP line is modeled as a random effect.

| Condition | Source | DF | Mean Sq | F Value | *p*-value |
|---|---|---|---|---|---|
| Control diet | L | 1 | 2.68 | 3.05 | 0.0821 |
| | $\epsilon$ | 241 | 0.88 | - | - |
| Restricted diet | L | 1 | 2.23 | 2.11 | 0.1479 |
| | $\epsilon$ | 231 | 1.06 | - | - |
| Both diets | L | 1 | 0.01 | 0.01 | 0.9175 |
| | D | 1 | 13.11 | 13.55 | 0.0003 |
| | L×D | 1 | 4.90 | 5.07 | 0.0248 |
| | $\epsilon$ | 472 | 0.97 | - | - |
